# Neck muscle spindle noise biases reaches in a multi-sensory integration task

**DOI:** 10.1101/182907

**Authors:** Parisa Abedi Khoozani, Gunnar Blohm

**Affiliations:** Centre for Neuroscience Studies, Queen’s University, Kingston, Ontario, Canada; Canadian Action and Perception Network (CAPnet), Toronto, Ontario, Canada; Association for Canadian Neuroinformatics and Computational Neuroscience (CNCN), Kingston, Ontario, Canada

## Abstract

Reference frame Transformations (RFTs) are crucial components of sensorimotor transformations in the brain. Stochasticity in RFTs has been suggested to add noise to the transformed signal due to variability in transformation parameter estimates (e.g. angle) as well as the stochastic nature of computations in spiking networks of neurons. Here, we varied the RFT angle together with the associated variability and evaluated the behavioral impact in a reaching task that required variability-dependent visual-proprioceptive multi-sensory integration. Crucially, reaches were performed with the head either straight or rolled 30deg to either shoulder and we also applied neck loads of 0 or 1.8kg (left or right) in a 3×3 design, resulting in different combinations of estimated head roll angle magnitude and variance required in RFTs. A novel 3D stochastic model of multi-sensory integration across reference frames was fitted to the data and captured our main behavioral findings: (1) neck load biased head angle estimation across all head roll orientations resulting in systematic shifts in reach errors; (2) Increased neck muscle tone led to increased reach variability, due to signal-dependent noise; (3) both head roll and neck load created larger angular errors in reaches to visual targets away from the body compared to reaches toward the body. These results show that noise in muscle spindles and stochasticity in general have a tangible effect on RFTs underlying reach planning. Since RFTs are omnipresent in the brain, our results could have implication for processes as diverse as motor control, decision making, posture / balance control, and perception.

**New & Noteworthy:** We show that increasing neck muscle tone systematically biases reach movements. A novel 3D multisensory integration across reference frames model captures the data well and provides evidence that the brain must have online knowledge of full body geometry together with the associated variability to accurately plan reach movements.

## Introduction

Different sensory and motor signals are encoded in different coordinates in the brain, e.g. early vision in eye/gaze-centered, primary arm proprioception in shoulder-centered. Conversions between reference frames are vital to transform signals into reference frames that are appropriate for processes as diverse as motor control, decision making, posture / balance control, and perception (Flanders et al., 1992; Buneo et al., 2002; Vetter et al., 1999; Blohm & Crawford, 2007). Previous studies have suggested that reference frame transformations (RFTs) should be regarded as stochastic processes which modulate the reliability of transformed signals (Alikhanian et al., 2015, Schlicht & Shrater, 2007; Burns & Blohm, 2010, 2011). Furthermore, several studies proposed that humans flexibly select the coordinates that minimize the effect of stochasticity (Sober & Sabes, 2005). Cue reliability-based multi-sensory integration studies have shown that stochastic RFTs affect human behavior (Schlicht & Shrater, 2007; Burns & Blohm, 2010, 2011); however, the sources of stochasticity in RFTs as well as the underlying mechanisms of how RFTs affect transformed signals remain unclear.

In order to accurately perform RFTs, the brain must have an estimate of 3D body articulation (Blohm & Crawford, 2007); i.e. an internal estimate of different body parts with regard to each other (such as eye re. head translation) as well as an estimate of joint angles (such as head/eye orientations). While the former is likely learned and does not change, the latter could stem from at least 2 sources, noisy afferent sensory signals (proprioception) and efferent copies of motor commands. Both signals are inherently variable due to the uncertainty of sensory reading and the variability of neuronal spiking (Poisson noise). Several studies have suggested that varying body articulation, e.g. the head roll angle, increases the behavioral variability due to signal-dependent sensory and neural noise affecting the RFT (Alikhanian et al., 2015; Schlicht & Shrater, 2007; Burns & Blohm, 2010, 2011). Signal-dependent sensory noise can arise from variability in the muscle spindle activity, the vestibular system, or both (Lechner-Steinleitner, 1987; Scott & Loeb, 1994; Cordo et al., 2002; Sadeghi et al., 2007; Faisal et al., 2008). Thus, larger joint angle estimates are accompanied by higher uncertainty (Wade & Curthoys, 1997; Van Beuzekom & Van Gisbergen, 2000; Blohm & Crawford, 2007), which results in an increased trial-to-trial variability in the RFT.

The effect of stochastic RFTs on the reliability of transformed signals has been studied using a multi-sensory integration task. Multisensory integration combines different sources of sensory information to create the best possible estimate of the state of our body within the environment in a way that is generally well captured by Bayes-optimal integration (Stein & Meredith, 1993; Landy et al., 1995; Atkins et al., 2001; Landy & Kojima, 2001; Kersten et al., 2004; Stein & Stanford, 2008; Ernst & Banks, 2002; Knill & Pouget, 2004). For instance, both visual and proprioceptive information can be combined in a reliability-weighted fashion to estimate hand position. It is believed (weak fusion hypothesis, Clark & Yuille, 1990) that prior to integration any signals must first be converted into a common coordinate system; this requires a (stochastic) RFT. Within this framework, the reliability of the transformed signal is affected by stochasticity in RFTs (Alikhanian et al., 2015), thus modulating the multisensory integration weights (Burns & Blohm, 2010; Burns et al., 2011). However, it is not clear how varying multisensory weights due to stochastic RFTs affects reaching movements to visual targets.

Here, we deployed a modified version of the standard visual-proprioceptive integration-based reaching task (Van Beers et al., 1999; Sober & Sabes, 2003, 2005) to systematically investigate the behavioral consequences of biases and variability in sensory estimates used for stochastic RFTs. We asked human participants to perform a center-out reaching task while the seen and actual hand positions were dissociated. In addition, reaches were performed with the head either straight or rolled 30deg to either shoulder and we also applied neck loads of 0 or 1.8kg (left or right) in a 3×3 design. Our results demonstrate that applying the neck load increased the variability of reach movements and biased the reaching behavior toward the applied load in all head roll orientations. Our prediction was that these effects on reaching behavior can be explained by a change in multisensory integration weights due to stochastic RFTs, which consequently enabled us to quantify the relative contribution of neck muscle spindles to the estimation of head roll angle. To test this hypothesis, we implemented a novel 3D stochastic model of multisensory integration across reference frames. Our model was able to capture the pattern of behavioral data well and allowed us to make two main conclusions: the effect of neck load on reaching behavior can be explained by changes in multisensory weights due to stochastic RFTs and the source of this stochasticity in RFTs is signal-dependent noise.

## Material and Method

### Participants

Nine healthy humans (8 male) between 20 to 35 years of age with normal or corrected to normal vision participated in our reaching task. They performed their reaching with their dominant right hand. Experimental conditions were approved by the Queen’s University General Board of Ethics and all the participants gave their written consent. Monetary compensation was provided for participating in the experiment ($10/hour).

### Apparatus

A virtual reality robotic setup (KINARM End Point Robot, BKIN Technologies) was used for performing the center-out reaching task. Participants stood in front of the robot while positioning their head by resting the forehead on the robot in front of the screen and their chin on a chinrest. Participants grasped a vertical handle attached to the robotic arm in order to reach to the viewed target on the mirrored surface. The vision of participants’ hand was occluded using an opaque board and eye movements were tracked using embedded technology (Eyelink 1000, SR Research). A pulley system and a helmet were used for measuring the head roll and loading the neck (see Figure 1 A and C).

**Figure 1.**
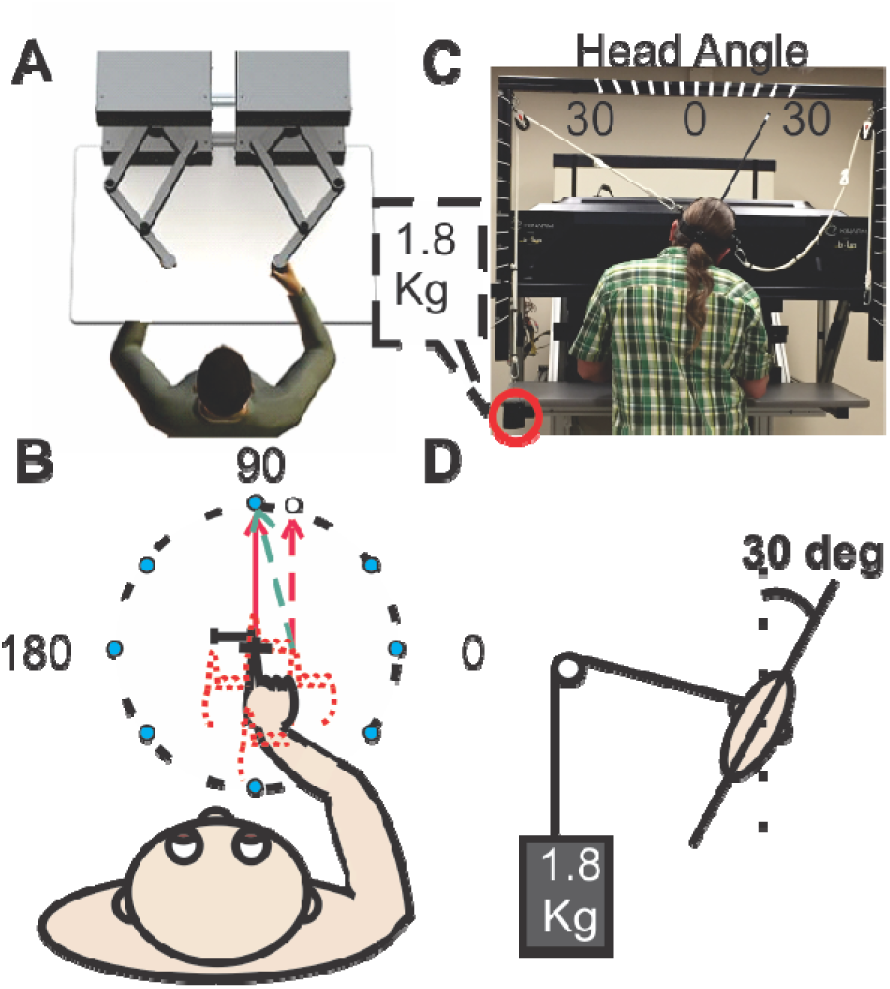
Apparatus-. A) KINARM end point robot (BKIN technology website) arrangement. B) Visual targets were distributed evenly on a 10cm-radius circle. The hand was shifted 2.5cm either vertically or horizontally while the visual indicator stayed at the center. C) Picture of the pulley system for measuring the head roll and loading the neck, in this picture the participant had 30CW HR and neck load on the left side. The attached indicator on the helmet was used to measure the head angle.

### Task Design

Participants stood in front of the robot and grasped the handle. At the beginning of each trial, participants were instructed to position their hand on the start position (cross) in the center of the display field. The robotic arm moved the hand toward the center and released it when the hand was within 3 centimeter of the central cross; a red dot representing hand position appeared at this point. After the participant positioned the hand correctly on the cross, one of the eight targets, distributed evenly on the circle with radius 10 cm, appeared. Participants were instructed to move through the target quickly and accurately while keeping their gaze fixated on the center cross. Once the participant’s hand begun to move (85 mm/s velocity threshold), the hand cursor disappeared. If they reached the target in less than 750ms, the trial was successful and participants would hear a successes beep, otherwise a failure beep was played indicating that the trial had been aborted and would have to be repeated. At the end of each trial,the center cross disappeared and participants had to wait 500ms to start the next trial. The next trial started with the reappearance of the center cross and the movement of the robotic arm driving the participant’s hand to the start position. This was to ensure that participants did not have visual feedback of their previous trial’s performance.

There were several different conditions in our experiment: The hand was physically shifted randomly either up/down or left/right with respect to the visual feedback of the hand. For example, participants would align their hand cursor to the center cross while their actual hand position was 2.5cm left of the cross. This discrepancy was introduced to enable us to measure the relative weighting of vision and proprioception in the multisensory integration process, similar to the logic employed in Sober and Sabes (2003, 5005) and Burns and Blohm (2010). In addition, the reaching movements were performed while the participants either kept their head straight or rolled their head 30deg toward each shoulder and while a neck load (0 or 1.8kg) was applied to the left or right side (the value of the weight was chosen to stimulate the same force as a 30deg head roll on neck muscles). Combinations of different head roll (HR) and neck load (NL) conditions are shown in Figure 2. We hypothesized that altering head roll and neck muscle force would create a conflict for head roll estimation as well as changing the signal-dependent noise which will affect the weights of multi-sensory integration. Participants completed 640 trials (5 hand positions * 8 targets * 16 repetitions) for each of the 9 combinations of head roll/neck load, for a total of 5760 trials (640*9) in 6 one hour sessions. In order to avoid any biases due to a specific order of experiment conditions, we employed Latin squares method to counter balance among different experimental conditions (Jacobson & Matthews, 1996).

**Figure 2.**
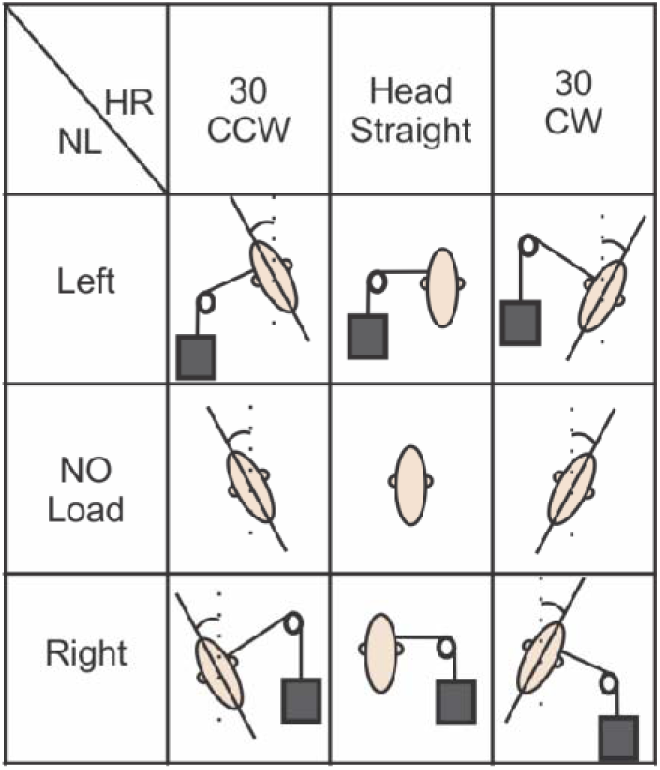
Experimental conditions. Participants performed the reaching task under 9 different combinations of HR and NL conditions during our experiment.

### Data Analysis

Hand and eye movement were captured with sampling rates of 1000Hz and 500Hz respectively. MATLAB software was used for offline analysis: A low-pass filter (autoregressive forward-backward filter, cutoff frequency = 50 Hz) was used to smooth the acquired data. First and second derivative of hand position data was calculated (using a central difference algorithm) to obtain hand velocity and acceleration. Trials in which participants moved their eyes after the visual target is displayed or moved their hand in a predictive direction except the target direction were removed (3% of overall trials). The time span from when participants started to move until their hand crossed a 9cm circle is defined as the initial movement duration. Movements were typically straight and had very little curvature; thus movement angle was derived through regression of data points acquired throughout the initial movement duration. Since the visual and proprioceptive hand position was dissociated, we defined visual movement as the movement obtained when subtracting visual hand from target information (red arrow, Figure 1B) and proprioceptive movement as the movement direction obtained when subtracting proprioceptive hand position from the visual target information (green arrow, Figure 1B). Subtracting predicted visual (proprioceptive) movement from the measured movement angle yielded the directional visual (proprioceptive) movement errors, which we used for our analysis. We then used an analytical model to capture the pattern of movement errors measured across conditions and targets (see model description below).

### Statistical Analysis

An n-way repeated measure ANOVA (rm-ANOVA) was used to assess the statistical differences (MATLAB 2013a, anovan.m) and post-Hoc analysis using the Bonferroni criterion (MATLAB 2013a, multcompare.m) was performed to assess the interaction between different parameters. A paired t-test (MATLAB 2013a, ttest.m) was used to assess the statistical significance in reach error variability for different head roll and neck load conditions. In all the statistical analysis p < 0.001 was considered as the criterion for statistical significance.

### Model description

The goal of our model was to understand which intrinsic and extrinsic variables were required to perform the RFTs accurately and more importantly, how variation of such variables affects human movement behavior. In order to understand the effect of RFTs on reach planning, we first explain the required steps in our model to plan a reach movement. Sober and Sabes (2003) proposed a two-step model for planning a reach movement in which first a movement plan is calculated by subtracting the hand position from the target position. Then this movement plan transformed to a desired change in arm angles through performing inverse kinematics. We extended previous models (Sober & Sabes, 2003; Burns & Blohm, 2010) that considered two steps for planning a reach movement: 1) calculating the movement plan and 2) generating the motor command. Several neurophysiology studies suggested that the movement plan is coded in visual (retinal) coordinates (Andersen & Buneo, 2002; Batista et al. 1999) while motor commands are coded in joint coordinates (Crawford et al. 2004). Following the same logic, in our model the two steps were performed in two different coordinates respectively: visual and proprioceptive coordinates. Visual information of hand and target positions were coded as retinal information in gaze-centered coordinates, X_h_=(x_1,h_,x_2,h_) and X_t_=(x_1,t_,x_2,t_) respectively (left panel in **Error! Reference source not found.** Figure 3), while the proprioceptive information of initial hand position was coded as joint angles in shoulder-centered coordinates, (**θ**_1,h_,**θ**_2,h_), (right panel in Figure 3**Error! Reference source not found.**).

**Figure 3.**
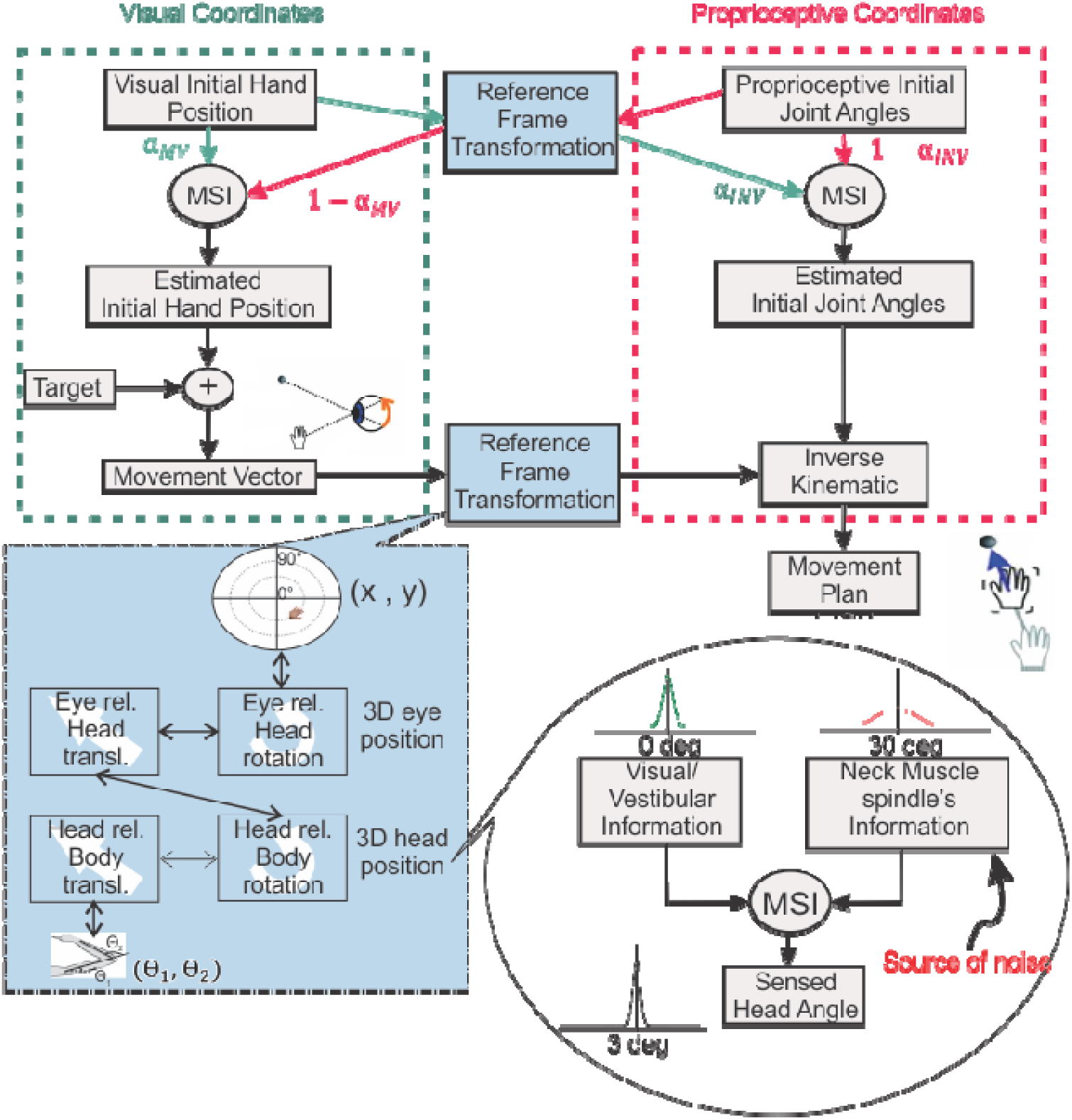
Model schematic. In order to perform the reach movement successfully, IHP is calculated in both visual and proprioceptive coordinates. In visual coordinate, IHP is computed by transforming proprioceptive information into visual coordinates. Visual and transformed proprioceptive information are weighted and combined based on Bayesian theory. A movement vector is calculated by comparing the estimated IHP and target positions. The same process takes place in proprioceptive coordinate to generate a proprioceptive IHP estimate. Using inverse kinematic, the transformed movement vector and IHP can be combined to calculate the movement plan based on the required changes in joint angles. The blue box represents the RFTs process. RFTs are performed by considering eye and head orientation as well as the translations between rotation centers of the body. The head orientation is estimated by combining visual/vestibular and neck muscle spindle information using Bayesian statistics (see Methods for details).

### Reference frame transformation (Blue box Figure 3)

In order to accurately transform information between the visual and proprioceptive coordinates the full body geometry must be taken into account (Blohm & Crawford 2007). This is specifically important when the head is not straight, i.e. rotating the head results in shifts of centers of rotation of the eye, head, and shoulder relative to each other (Henriques & Crawford, 2002; Henriques et al., 2003). To capture this, we performed a series of rotations (R) and translations (T), formulated in equations (1) and (2) respectively.

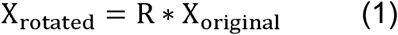

Where 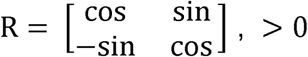 holds for clock wise rotations.

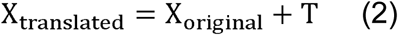

In the following section, we explain the required steps to transform hand position from eye-centered to shoulder-centered coordinates.

#### Retinal-to-shoulder transformation

As it is depicted in Figure 3, in order to transform retinal-coded information into joint-coded information the theoretically required sequential transformations can be done by first transforming retinal to head coordinates, then from head to shoulder and finally from should to joint coordinates (Note that this is likely different from how the brain is performing this transformation):

1. Retinal-to-head

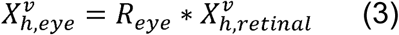

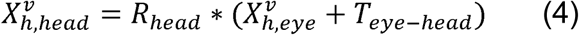 In which *R*_*eye*_ and *R*_*head*_ are rotations based on eye angle and head angle respectively and *T*_*eye-head*_ is the translation between eye and head which is the distance between the center of two eyes (eye-centered coordinate) and the joint of head and neck (head centered coordinate). 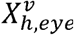 is the visual information of hand position in eye-centered coordinate: Subscript ‘h’ represents information related to the hand position and the following subscript represents the related coordinate at that step. In addition, we deployed superscripts ‘v’ or ‘p’ to dissociate if the information is originally provided by vision or proprioception respectively. All the following parameters have the same pattern.
2. Head-to-shoulder

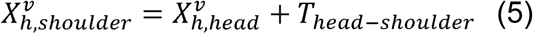 Since the body was upright, a translation is sufficient to perform the transformation between the shoulder and head. In our setup, the shoulder was located downward and to the right of the head.
3. Shoulder-to-joint

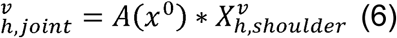 In which *A(x*^*^0^*^) is the forward kinematic matrix and has the same form as equation (7) by Burn and Blohm (2010), since our experimental configuration is the same. In order to transform the information from joint angle coordinates to retinal coordinates, the same procedure can be performed only in the reverse order (since we used the same configuration as Burns and Blohm (2010), both forward and inverse kinematic matrices have the same format).

In addition to the full body geometry, we considered the noise of transformation in our model. Similar to Burns and Blohm (2010), we have two noise component resulted from the transformation: fixed transformation noise 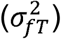 to simulate the fact that any transformation has a cost (Sober and Sabes 2005), and variable transformation noise 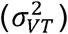 to simulate the different head orientations and neck load conditions of our experiment (this is the same as the variability in the estimated head angle).

### Estimating head angle

As mentioned in the previous section, participants performed reaching with different head roll and neck load conditions. Therefore, our model must include a head angle estimation component as a crucial part of the RFTs processes. Previous studies showed that humans combine visual, vestibular, and neck proprioceptive information for estimating head orientation, similar to a Bayesian optimal observer (Mergner et al., 1983, 1991, and 1997; Clemens et al., 2011; Alberts et al., 2016). For instance, Mergner et al. (1991) demonstrated that the stimulation of neck muscles by rotating the trunk on a fixed head caused a sensation of head rotation and also increased the uncertainty of head position estimation. In addition, two studies carried out in Medendorp’s group demonstrated that the noise in both vestibular and proprioceptive information should be considered signal-dependent (Clemens et al., 2011; Alberts et al. 2016). Therefore, we used a similar principle for our head angle estimation in RFTs processes. Thus, following the same rational, we included neck load in our experimental condition with the goal of investigating the contribution of the mentioned sources of information for estimating the head angle. Assuming that each source of information has a Gaussian distribution, the head angle signal has a Gaussian distribution as well and its mean and variance can be estimated as follows:

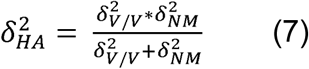

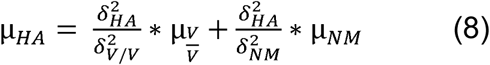

In which 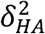, 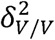 and 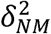 are associated variability in head angle estimation, visual/vestibular information, and neck muscle information respectively and *μ*_*HA*_*, μ*_*V/V*_, and *μNM*are the associated means in the same order. Therefore, we also were able to extract the relative visual/vestibular vs. neck muscle contribution in estimating head angle 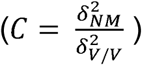

As mentioned earlier, one of the key features of our model is including signal dependent noise in our RFTs: The assumption is that when we roll the head, the variability of both vestibular and neck muscle spindle signals increase due to higher signal value. In addition, applying the neck load increases the force on the neck muscle which results in increasing the variability of neck muscle spindle signal. In the conditions of applying the neck load while the head is not straight, the two forces on the neck muscle are combined in order to drive the predicted neck muscle force. Therefore, we differentiated the variability for the head straight and no load condition from the other head roll and neck load conditions. Similar to Vingerhoets et al. (2008), we used a linear model to explain the increase in variability due to increase in the signal value:

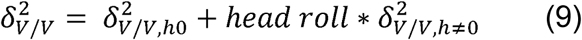

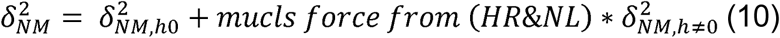

In which 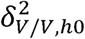 and 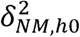 are visual/vestibular and neck muscle variability for head straight condition and 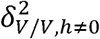 and 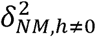 are the ones for other experimental conditions. This will result in having μ_*HA,h0*_ and μ_*HA,h≠ 0*_.

At the final step, the required head angle for the transformation (_*HA*_) is derived by scaling the estimated head angle (μ_*HA*_) (obtained by sampling from the above Gaussian distribution) by a gain factor β:_*HA*_ =β * μ_*HA*_.

### Multisensory integration

In order to estimate the initial hand position (IHP), visual (V) and proprioceptive (P) information are combined using multisensory integration principles. In our model, the multisensory integration is happening twice: once in visual coordinates (coded in Euclidean) in order to calculate the movement vector (MV) and once in proprioceptive coordinates (coded in joint angles) in order to generate the motor command using inverse kinematics (INV). We assumed that each piece of information has a Gaussian distribution (before and after RFTs) and therefore using multivariate Gaussian statistics the mean and covariance of the combined IHP estimated from vision (V) and proprioception (P) in each coordinate can be written as:

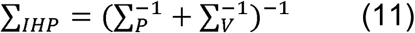

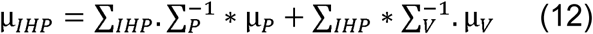

Where *∑*_*IHP*_ is the covariance matrix of IHP and *∑*_*V*_ and *∑*_*P*_ are covariance matrices of visual and proprioceptive information respectively. Similarly, μ_*IHP*_, μ_*P*_, and μ_*V*_ are the mean values (in the vector format) for IHP, visual, and proprioceptive information. Therefore, the visual weight in each of the visual and proprioceptive coordinates is calculated as:

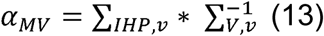

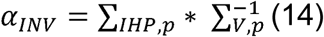

Where *α*_*MV*_is the multisensory integration weight for visual information in visual coordinates and *α*_*INV*_ is the multisensory weight for visual information in proprioceptive coordinates. Where *∑*_*IHP,v*_ is the covariance matrix of IHP in visual coordinates and *∑*_*V,v*_ is the covariance matrix of visual information in visual coordinates. Similarly, *∑*_*IHP,p*_ is the covariance matrix of IHP in proprioceptive coordinates and *∑*_*V,p*_ is the covariance matrix of visual information in proprioceptive coordinates.

### Final motor command and movement direction

After estimating the IHP, the desired movement vector is calculated by subtracting the hand position from the target positon;*Δx=tar-μ*_*IHP,v*_.We used the velocity command model (Sober & Sabes 2003; Burns & Blohm 2010) to transform the desired movement vector to the required motor command: 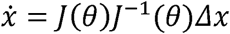(where J(θ)and J^−1^(θ)have the same form as equation (16) and (17) in Burns and Blohm 2010).

At the final step the movement direction is calculated by transforming the movement command from Euclidean coordinates to polar coordinates using the following equations:

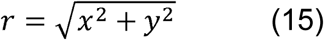

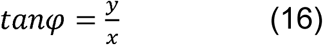

### Generating quantitative model predictions

In order to generate our model predictions we used a Monte Carlo approach (Binder & Heermann, 2002); we assumed that the sensory information (visual and proprioceptive information of initial hand position, visual/vestibular and proprioceptive information of head position) can be sampled from a Gaussian distribution with a specific mean and covariance matrix. Then, the RFT procedure is performed on each sample based on sampled head roll signals to obtain the distribution of the transformed signal. The movement direction was calculated for each sample and the final movement mean and variance were calculated based on this distribution. The model code is available on Github (https://github.com/Parisaabedi/NeckMuscleLoad).

### Model parameters

Based on average body physiology, upper arm and lower arm (including fist) lengths were set constant to L1 = 30 and L2 = 45 cm respectively. Shoulder location was assumed 30 cm backward from the target and 25 cm rightward of the target, the distance between eye and top of the head considered 13 cm, and the head length considered 28 cm (40 cm including the neck). IHPs and target positions were taken from the experimental data.

There were seven free parameters in the model, i.e. the variance of both proprioceptive 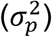 joint angles and visual IHP 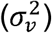 - we assumed that the two dimensions in both coordinates are independent with the same variability-, the visual/vestibular vs. neck muscle spindle contribution factor (C), the variance of head angle estimation for head straight 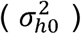, a fixed reference frame transformation cost 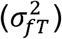, and a variable reference frame transformation cost.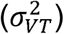.

As it is mentioned before, the 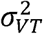 is resulted from the variability in the head angle estimation; 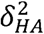 By substituting 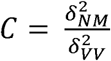 equation (7), we were able to extract the variance of neck muscle spindles 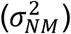 and visual/vestibular 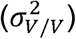 Furthermore, we added an additional variance component to account for the added variability during performing the planned movement 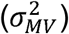.

In order to estimate the model parameters we used a standard maximum likelihood procedure. We calculated the negative log-likelihood of the angular reach error data to fit on the proposed model given parameter set *ρ* as:

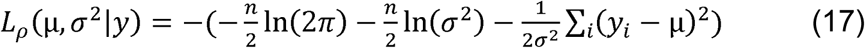

Where (μ,σ^2^) are the mean and variance driven from the model given the parameter set *p,n* is the number of data points and *yi* is each data point from the experiment. It should be noted that (μ,σ^2^) are calculated separately for each of the 360 experimental conditions: 8 visual targets * 5 IHPs * 3 head rolls * 3 neck loads. We then searched for the set of parameters which minimized the *L*_*p*_ over the parameter space using ‘fmincon.m’ in MATLAB 2017. Table 1 provides the fitting values for different model parameters for individual participants along with confidence interval for each parameter. We added one additional parameter C, which indicate the contribution of neck muscle information compared to combined visual/vestibular information by dividing the first by the second.

**Table 1.**
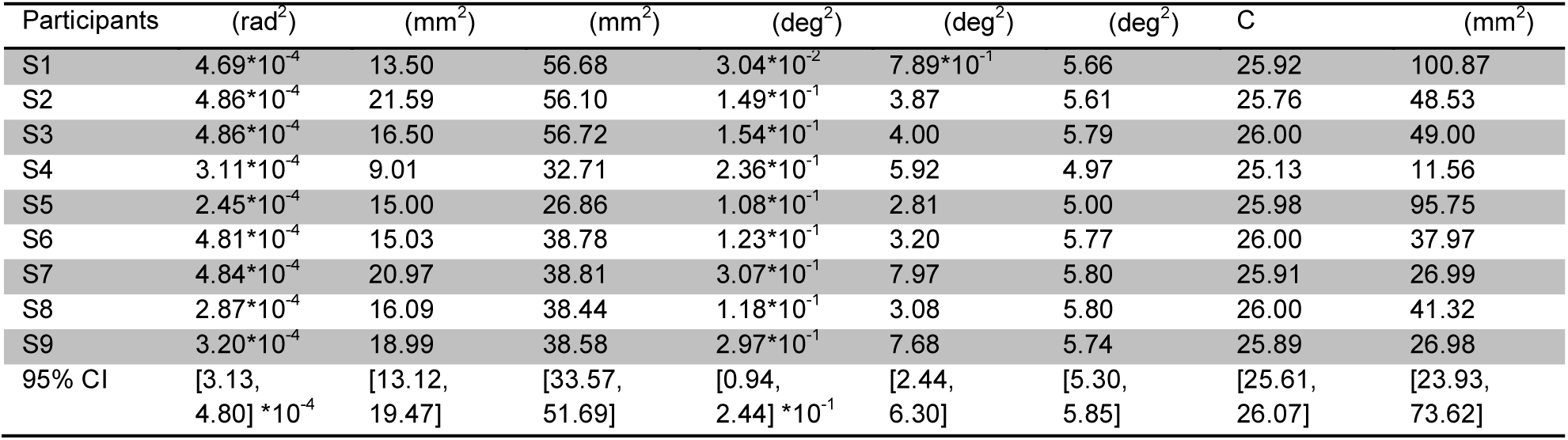
Model parameter fits.

## Results

Previous work (Burns & Blohm, 2010; Schlicht & Schrater, 2007; Sober & Sabes 2003) suggests human behavior is affected by stochastic RFTs. Burns and Blohm (2010) showed that rolling the head will increase the variability of reach movements and argued that could be due to the signal dependent noise in the sensed head angles: rolling the head increases the amplitude of the sensed head angle and the associated variability accordingly. Here, our goal was to investigate the sources of stochasticity in RFTs and the effect of such stochasticity on human reaching movements. To this aim, we asked human participants to perform reaching movements while their head was either straight or rolled toward each shoulder and a neck load of 0 or 1.8kg was applied to the right or left side in a 3×3 design. The experimental logic was that applying head roll and neck load will vary the sensed head angle and the associated noise due to signal-dependent noise. Since RFTs are based on these sensed angles, applying head roll / neck load increases the stochasticity of RFTs which modulates the multisensory integration weights and thus resulting in more variable and potentially biased reaching movements compared to the condition where the head is straight and no load is applied.

### General Observations

A total of 51840 trials were collected, with 1529 trials being excluded due to either eye movements or predictive hand trajectories. We used directional reach errors to determine how participants weighted their visual information vs. proprioceptive information. Directional reach error (in angular degrees) was computed by subtracting proprioceptive (visual) hand-target direction from overall movement direction (see Methods), where 0deg means no deviation from proprioceptive (visual) hand-target direction. By introducing the shift in the visual feedback of the initial hand position, a discrepancy between visual and proprioceptive information was created and as a result, we could determine how visual and proprioceptive information was weighted and integrated based on how participants responded to this discrepancy.

To evaluate how humans weight visual and proprioceptive information, we compared reach errors for each hand offset condition. In order to calculate the reach errors, we can use either the visual hand-target direction (red line in

**Figure 1B**) or the actual (proprioceptive) hand-target direction (green line in

**Figure 1B**). We called the first one visual reach errors and the second one proprioceptive reach errors and used them for different sections of this manuscript in order to show the effects more clearly. The difference in reach errors among different hand offsets indicates that both visual and proprioceptive information were used during reach planning. Figure 4 displays both proprioceptive and visual reach error curves across target directions for different initial hand position conditions for head straight and no load condition.

**Figure 4.**
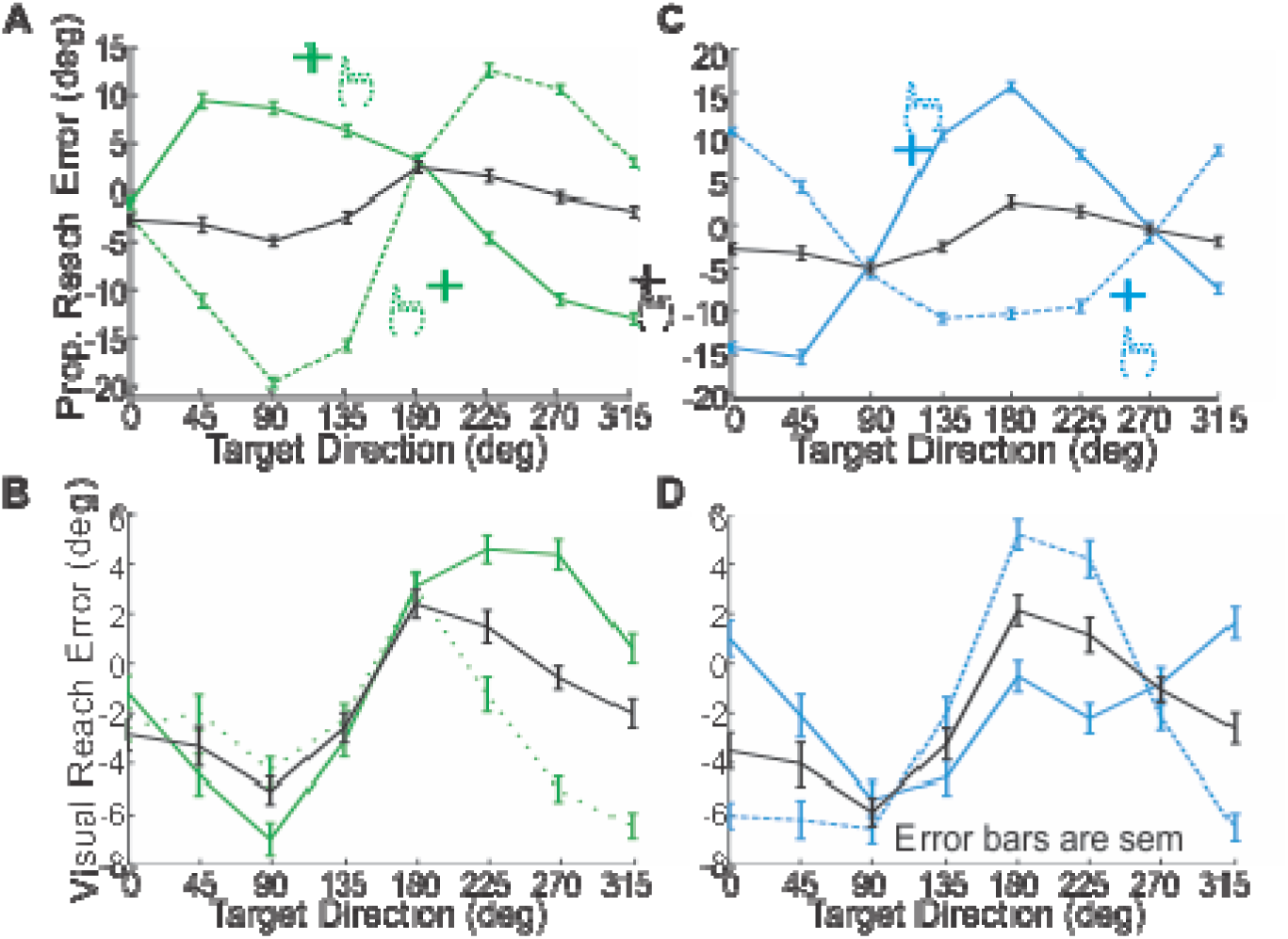
Reach error curves. Reach errors are calculated for each target by subtracting the proprioceptive or visual hand-target direction from the performed reach movement. Solid colored lines are representing upward/rightward shifts. A,C) proprioceptive reach error curves: (A) reach errors for horizontal hand shift (green, stays the same in the rest of the manuscript) and (C) reach errors for vertical hand shift (blue, stays the same in the rest of the manuscript). B,D) visual reach error curves: (B) reach errors for horizontal shift and (D) reach errors for vertical shifts.

To quantify these weights, we fitted a previously proposed model (Sober & Sabes 2003) to the normalized data. The data was normalized by subtracting the 0 hand offset from the other hand offsets. The model by Sober and Sabes (2003) fits our data well (Figure 5, R-squared for pooled data across all participants was equal to 0.91 and 0.93 respectively for the right and left panels) and confirms that the participants used both visual and proprioceptive information to plan their reach movement. Based on this close fit of our data to the model, we can now use this model in a first step to investigate how head roll and neck load affects the weighting of vision and proprioceptive information about the hand.

**Figure 5.**
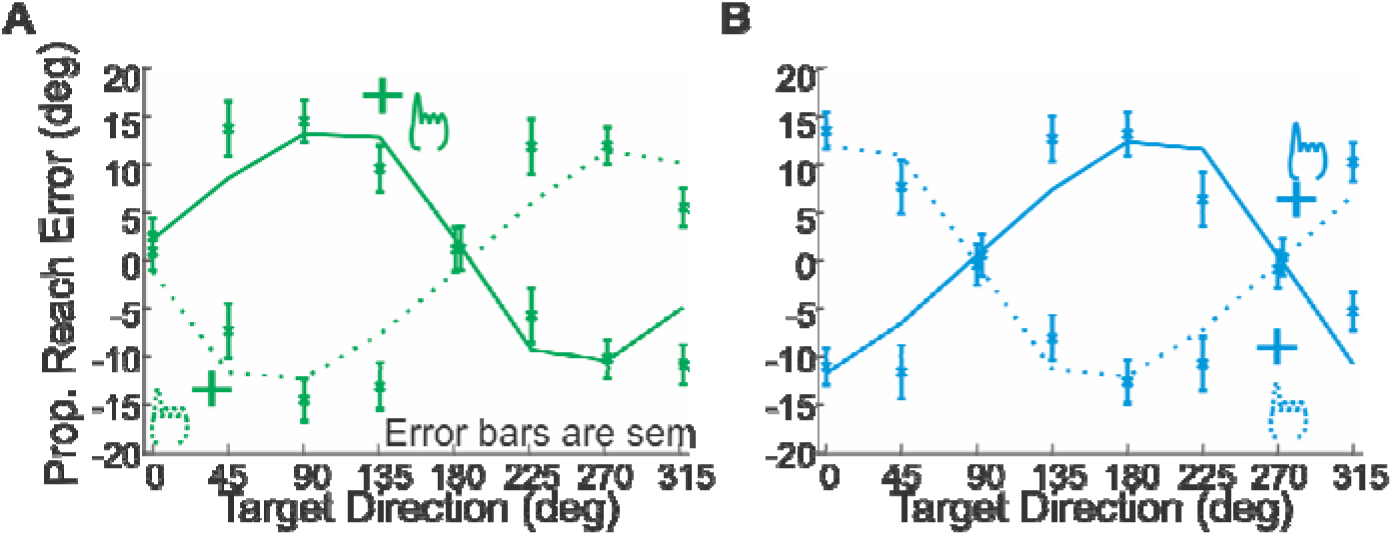
Sober and Sabes 2005 model fit on the data. The reach error curves are normalized to zero by subtracting the 0 hand offset from the other hand offsets.

### Head Roll effect

Participants performed the reach-out movements for different head roll conditions: 30deg counter clock wise (CCW), 0deg, and 30deg clock wise (CW) head roll. In the first step we examined if the same effect reported by Burns and Blohm (2010) could be reproduced. As the author explained changing the head roll had two different effects on the reach error trajectories. First, the reach error curves shifted up/down-ward and second the variability of reach errors increased for the tilted head conditions compared to the head upright condition.

Figure 6 depicts the effect of changing HR on both reach errors and movement variability. As it can be seen, there are both a bias effect and an increased variability effect for altering the head orientation compared to head straight. The n-way ANOVA with factors HR, target directions, and participants showed a significant main effect for altering head orientation, F(2,98) = 11.85, p< 0.01; and significant interaction between reaching to different targets and different HR conditions, F(14,98) = 5.59, p < 0.01; which shows that the effect of altering HR is different for different target directions. Bonferroni-corrected post-hoc analyses indicated that the bias effect was significant among all the HR conditions. Regarding movement variability, we performed a paired t-test across all participants for each HR condition vs. no HR condition: The increase in standard deviation due to the rolled head is significant for both sides, HR = 30deg CW vs. HR = 0: t(8) = −3.6133, p < 0.01; HR = 30deg CCW vs. HR = 0: t(8) = −5.6011, p < 0.01. These results are consistent with the results reported by Burns and Blohm (2010).

**Figure 6.**
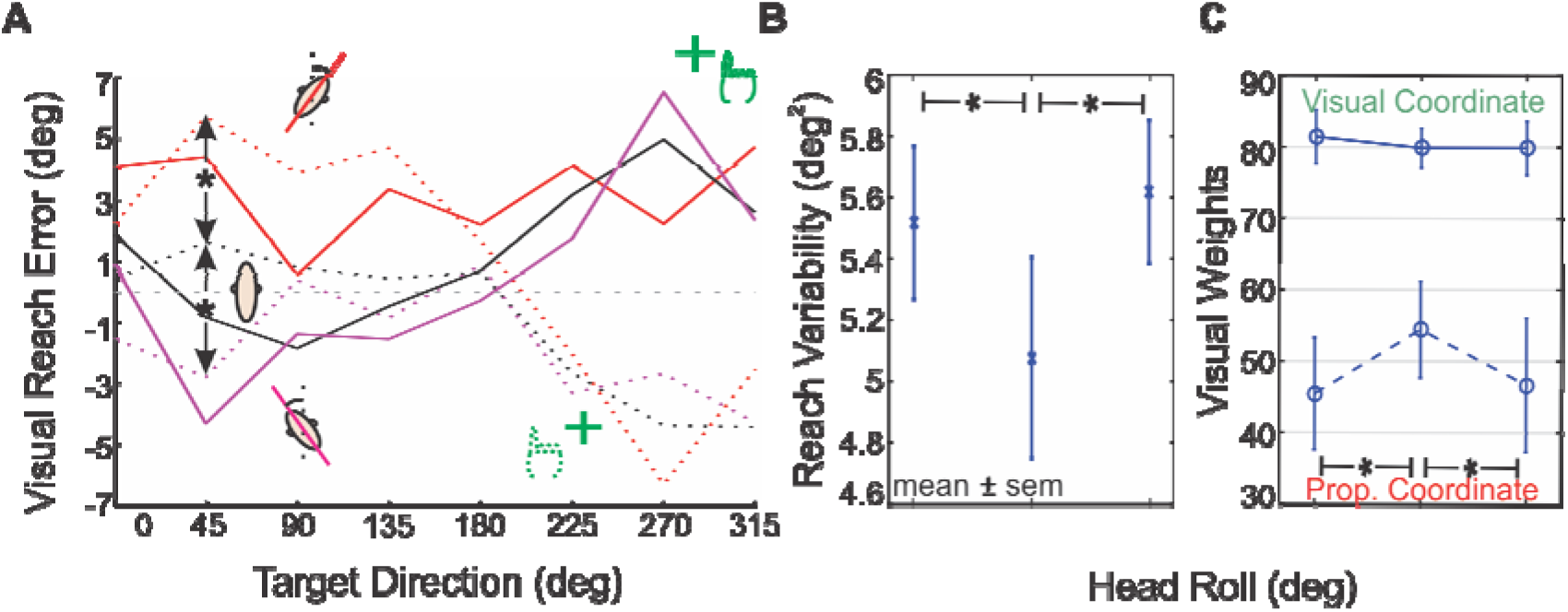
Effect of varying head roll on reach movement behavior. A) Reach error curves (solid line for IHP shifts to right and dotted line for IHP shifts to left) shifted upward for CW head roll and downward for CCW head roll compared to the head upright condition (n-way ANOVA, F(2,98) = 11.85, p < 0.01). B) The movement variability increased significantly for rolled head conditions compared to the head upright condition (paired t-test, p < 0.01). C) Visual weights derived by fitting the Sober and Sabes (2003) model on the data. We didn’t find any significant change in visual weights in visual coordinate for different head roll conditions, while the visual weights significantly decreased in proprioceptive coordinates. Significance was tested using paired t-test (P < 0.05 is considered as a significant difference).

We also used the Sober and Sabes (2003) model to extract the weights for different conditions. There, the visual and proprioceptive weights were the two free parameters of the model (Sober & Sabes, 2003) which is used to estimate the hand position, by integrating visual and proprioceptive information, in two different stages: visual weight in the movement planning stage and proprioceptive weight in the motor command generating stage. Therefore, the weights can be extracted after fitting the model on the data. As it is depicted in Figure 6, the visual weights in visual coordinates did not change very much by varying head roll, however, the visual weight in proprioceptive coordinate decreased for rolled head conditions. This is consistent with our hypothesis that higher noise in RFTs results in lower reliability of transformed signals which leads to higher weights for proprioceptive information in the proprioceptive coordinates compared to the head straight condition.

### Neck Load effect

In addition to altering the head roll a neck load (rightward, no load, or leftward) was applied. We assumed that if the neck load was not taken into account, there should be no difference in the reach errors between the neck load conditions and no load condition. Alternatively, if the neck load was taken into account in estimating head roll, then we expected to observe similar effects as during head roll; up/down-ward shifts in reach error curves and increased movement variability. This is because loading the neck while the head is upright would create a discrepancy between the neck muscle spindle information and the combined visual/proprioceptive information. In addition, due to signal dependent noise, the neck muscle information should become less reliable when the neck is loaded compared to the no load condition. Consequently, the sensed head angle estimated by integrating neck muscle and visual/vestibular signal should be biased toward the neck load and have more variability resulting in biased and more variable movements.

As it can be seen in Figure 7(A), applying the neck load created an up/down-ward shift of the reach movement error curves. An n-way ANOVA with factors NL, target location, and participants revealed a significant main effect for different NL, F(2,98) = 6.12, p < 0.01. Bonferroni-corrected post-hoc analyses indicated that the bias effect was significant among all the NL conditions. The interaction between targets and different NL was not significant, F(14,98) = 1.06, p = 0.402, which means that the effect of varying NL on reach movement was independent of different target directions.

**Figure 7.**
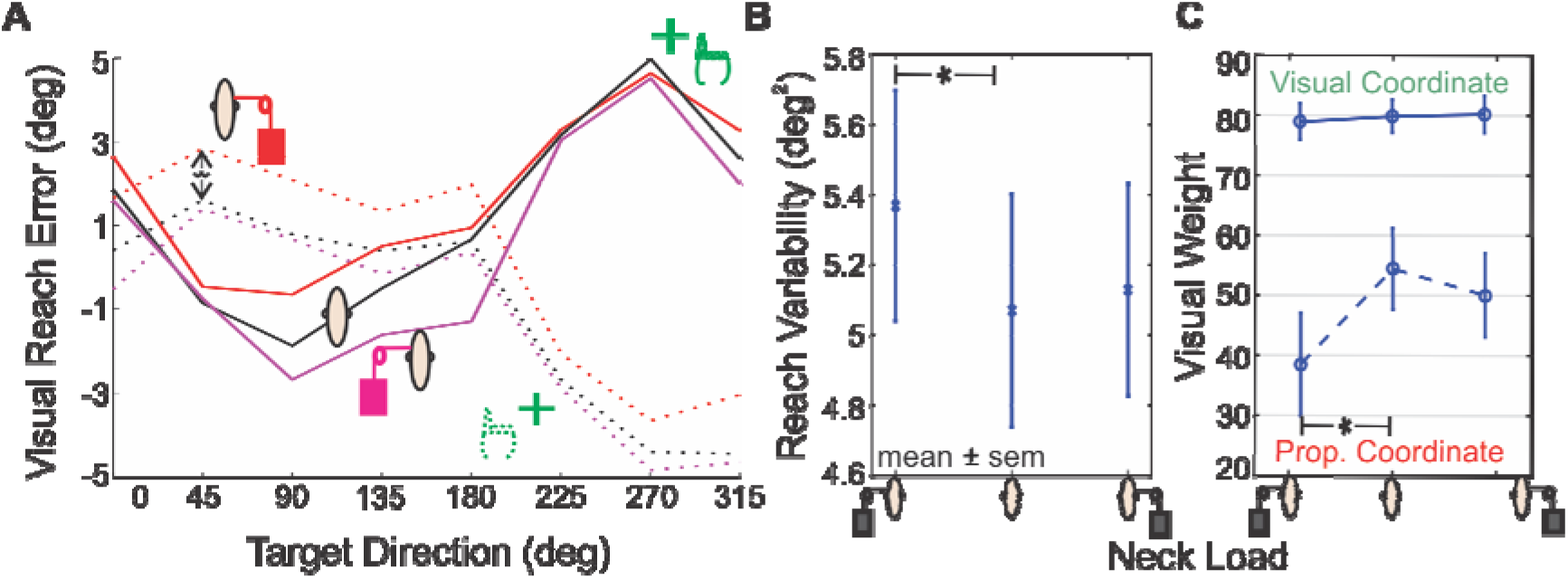
Effect of applying neck load on reach movement behavior. A) Reach error curves (solid line for IHP shifts to right and dotted line for IHP shifts to left) are shifted upward for applying neck load on the right (n-way ANOVA, F(2,98) = 6.12, p < 0.01). The shift in reach error curves for applying neck load on left is not statistically significant. B) The movement variability is increased significantly for applying the load on the left compared to the no load condition (paired t-test, t(8) = 2.7552, p = 0.0283). C) The visual weights derived by fitting the Sober and Sabes (2003) model on the data. We only observed a significant change in visual weight in proprioceptive coordinate due to applying neck load on the left side. Significance was tested using paired t-test (P<0.05 is considered as a significant difference).

Figure 7(B) represents the variability of reach errors in the no load condition vs. neck loaded conditions. As the figure demonstrates, the variability of reach errors is higher for applying the load compared to no load condition. We performed paired t-tests between all three different conditions across all eight participants. Movement variability was significantly higher for applying the load on the left side compared to no load condition t(8) = 2.7552, p = 0.0283. The paired t-tests revealed no significance difference among other conditions.

### Comparison

So far, we showed that there are both biases and increased movement variability effects for either applying NL or HR. In the next step, we compared the variability of reach movements in the NL conditions vs. HR conditions. Based on stochasticity in RFTs we expected to have higher variability for higher amplitudes of head angle during different experimental conditions. For example, we predicted to have higher movement variability for applying only HR compare to applying only NL or have the highest variability for conditions in which both HR and NL are applied in the same direction.

Based on Figure 8 the variability in HR conditions is higher than in NL conditions. We first ran paired t-tests between each condition separately and the significant statistical differences are shown in the Figure 8. Applying the load on the left side increased the variability compared to the control condition. Then, we performed paired t-test between combined similar conditions: for example both head upright and neck load on either side are combined and created the overall NL condition. The paired t-test between HR condition and NL condition showed a significant difference, t(8) = 2.7444, p = 0.0287; the difference between HR condition and control condition was significant as well, t(8) = 2.7444, p = 0.0020; however the difference between the control and NL conditions was not significant. Together all of the above observations provide evidence for the existence of signal dependent noise in the head angle estimation and consequently the RFTs processes. However, it is not clear how such stochastic RFTs affect the reaching movement. First, contrary to the initial hypothesis no modulation of variability was observed by varying NL while the head was rolled CW/CCW. In addition, in all the conditions we observed larger effects when rolling the head on reach errors for targets away from body (45-135 degree) compared to the targets toward the body (215-315 degree). Both previous models (Sober & Sabes 2003 and Burns & Blohm 2010) fail to explain the previousely mentioned effects. Based on both previous models, there shouldn’t be any difference in biases effect due to head roll condition and they predict a constant up/down –ward shift in the reaching error curves. We propose that these effects can be explained by a Bayesian framework which performs geometrically accurate RFTs.

**Figure 8.**
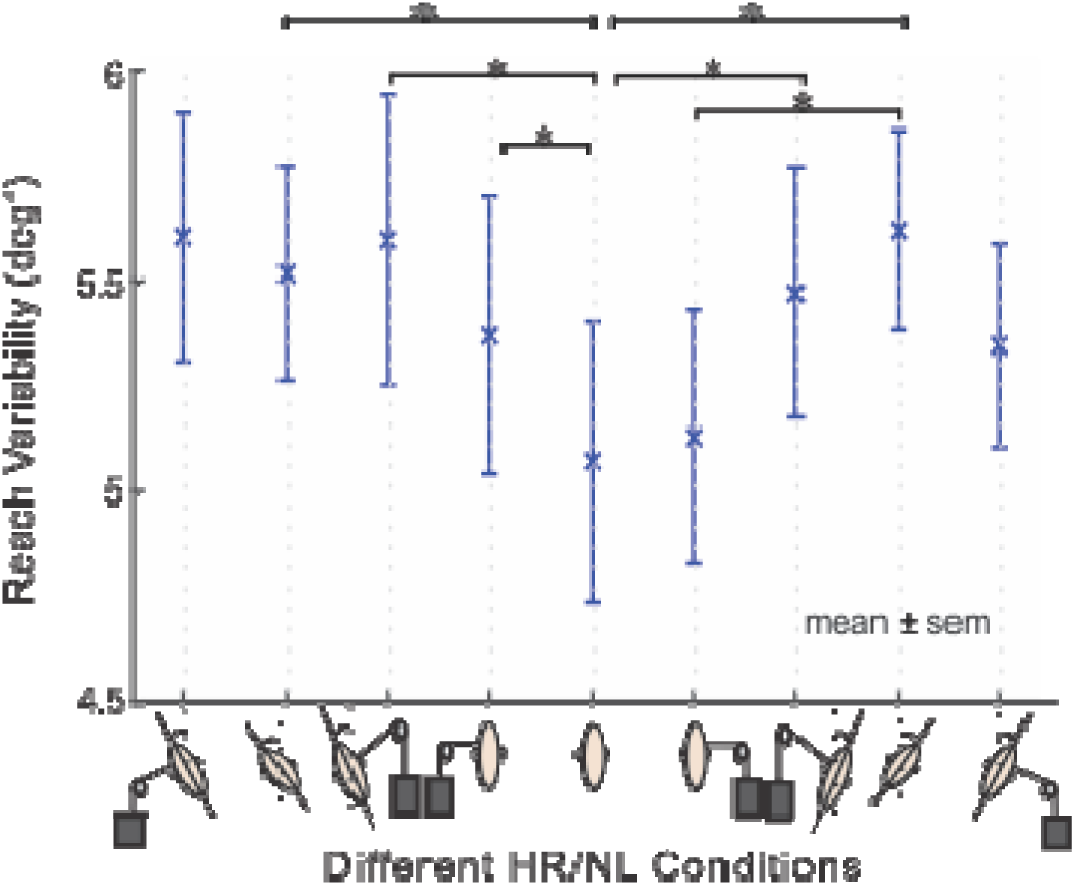
Effect of different experimental conditions on reaching movement variability. Head upright and no load condition (considered as the control condition) and the combined HR/NL conditions are sorted based on the expected increase in the variability based on the signal-dependent noise hypothesis right and left of the control condition. Rolling the head consistently increased the variability compared to the control condition. Significance was tested using paired t-test (P<0.05 is considered as a significant difference).

### Modeling the stochastic reference frame transformations

The above analyses demonstrate that RFTs should be considered as stochastic process. Therefore, to understand the effect of such stochastic RFTs on reach planning we developed a Bayesian model of multi-sensory integration for reach planning and explicitly included the RFTs.

**Error! Reference source not found.** Figure 3 depicts the schematic of our proposed model. The working principles of our model are similar to previous ones (Sober & Sabes, 2003; Burns & Blohm, 2010) with the addition of an explicit head orientation estimator (Figure 9, blue box). In summary, our model calculates the required reach movement through first calculating the movement vector in visual coordinates, by comparing estimated initial hand position and target position, and then generates the movement commands by transforming the movement vector from visual coordinates to proprioceptive coordinates.

**Figure 9.**
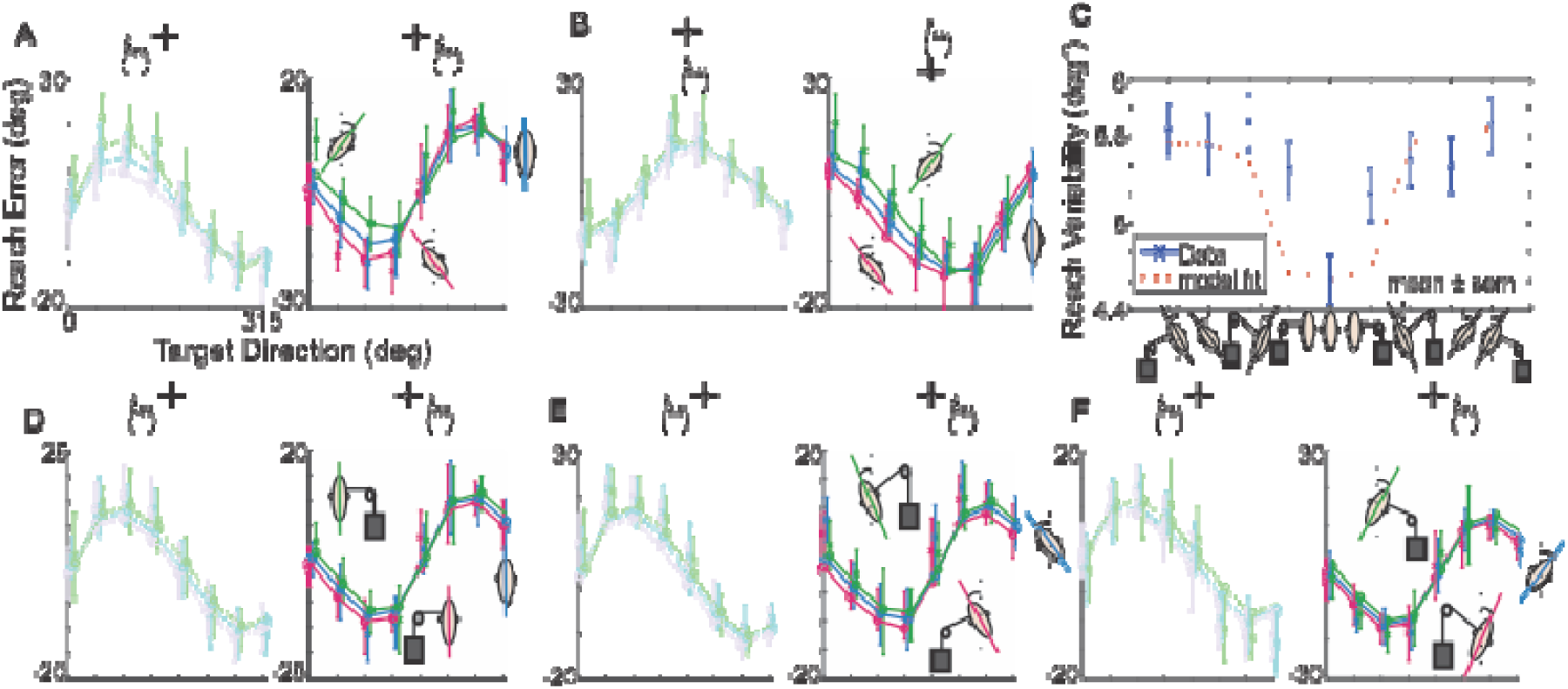
Model fit for a sample participant (#6). Model fit on the reach error curves for different IHPs and HR/NL conditions. A-B) model fit on the reach error curves for varying head orientation without applying neck load: A) solid line represents results for IHP shifts to the right and dotted line represents results for IHP shifts to the left, B) solid line represents results for IHP shifts to the up and dotted line represents results for IHP shifts to the down. C) model fit on the changes in movement variability due to varying HR and NL conditions. D-F) Model fit on reach errors for varying NL for different HR conditions. Only data for horizontal shifts are presented. The results for vertical hand shifts are similar.

We added several crucial features to the proposed model compared to the previous models (Sober & Sabes 2003, 2005; Burns & Blohm 2010). First, we explicitly included the RFTs. The RFTs processes transforms information between different coordinates considering the full body geometry; head orientation, eye orientation, head-eye translation, and head-shoulder translation. In addition, to perform the required transformations, we included a head angle estimator. The head angle estimator combines muscle spindle information and visual/vestibular information in a statistically optimal manner. Similar to Burns and Blohm (2010), we modeled both mean behavior and the associated variability for each source of information; vision, proprioceptive, vestibular, and muscle spindles. To examine the effect of noisy transformations on the visual/proprioceptive information, we deployed Monte Carlo simulations. This method gave us the opportunity to explicitly study the effect of RFTs on the covariance matrices and consequently the MSI weights.

### Model fit

In the following section we first provide the fitting results for a sample participant (#6) and then evaluate the fitting results across all nine participants. Figure 9 provides the fitting vs. data for participant # 6. Figure 9 (A and B) depicts model fitting for all different initial hand positions for different head rolls while no neck load was applied. As it can be seen, our model is able to accurately capture the reach errors for different IHP and HR conditions. Figure 9C provides the model prediction for changes in variance for different conditions. Error bars were derived using bootstrapping with N=1000. Since the results for horizontal and vertical hand shifts are very similar, for all the other conditions we only provided the results for the horizontal initial hand shifts. Figure 9D-F depicts the fitting for varying the NL for different Head angles; 0°, ±30°.

After demonstrating that our model was capable of predicting the reach error behavior for a single participant, Table 1 summarizes the fitting results for all the participants. The most interesting finding here is the relatively higher contribution of visual/vestibular signal compared to neck muscle spindle (C≈26). This is was consistent across all the subjects. We also observed very high movement variability across our participants.

Figure 10 provides the model prediction vs. data for both reach errors and variances for different experimental conditions. Different participants are differentiated by different colors. We used several different analyses to evaluate the goodness of our model fit. First, we calculated r-squared value for each individual participants and the pool of all the participants: [54, 61, 56, 75, 50, 60, 66, 71, 71, and 94] for S1-S9 and the pool of data respectively. Secondly, since the variance data was very noisy, we grouped them in bins and calculated the confidence interval for each predicted variance using the following equation (J. S. Williams, 1962):

**Figure 10.**
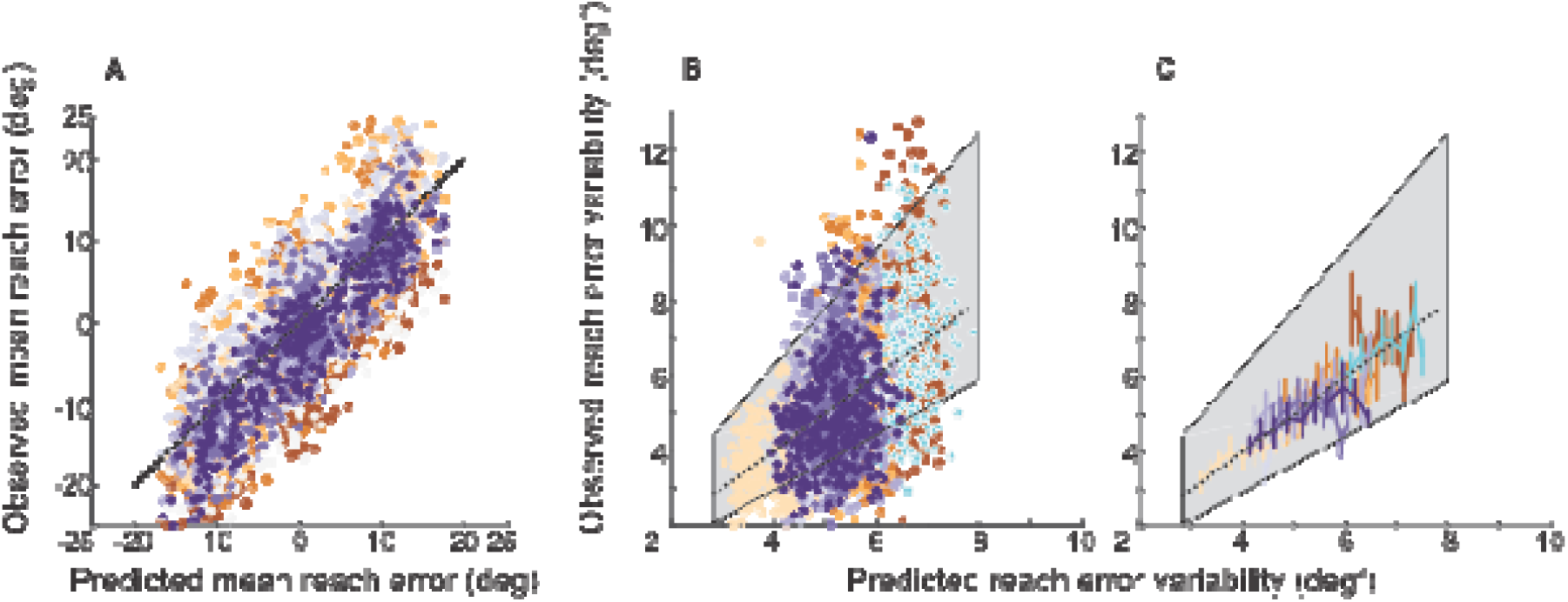
Model predication vs. observed data for each individual participant. Data for each individual participant was fitted to our model. Each color represents an individual participant. A) The model prediction vs. observed data for reach errors. B) The model prediction vs. observed data for reach variability. C) Same data as in section B grouped into bins of 0.25 deg2 (mean and standard error). The gray box represents the confidence interval for predicted variances based on our model.

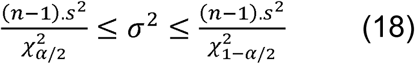

In which σ^2^ is the population variance,*S*^2^ is the sample variance,*n* is the sample size, and 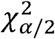 is chi-square distribution. Since we wanted to find the 95% confidence interval, we set α=0.05 The boxed colored area in Figure 11(B, C)is the calculated confidence interval for the variances. Based on this analysis, we could see that our model provides a decent fit on the data.

**Figure 11.**
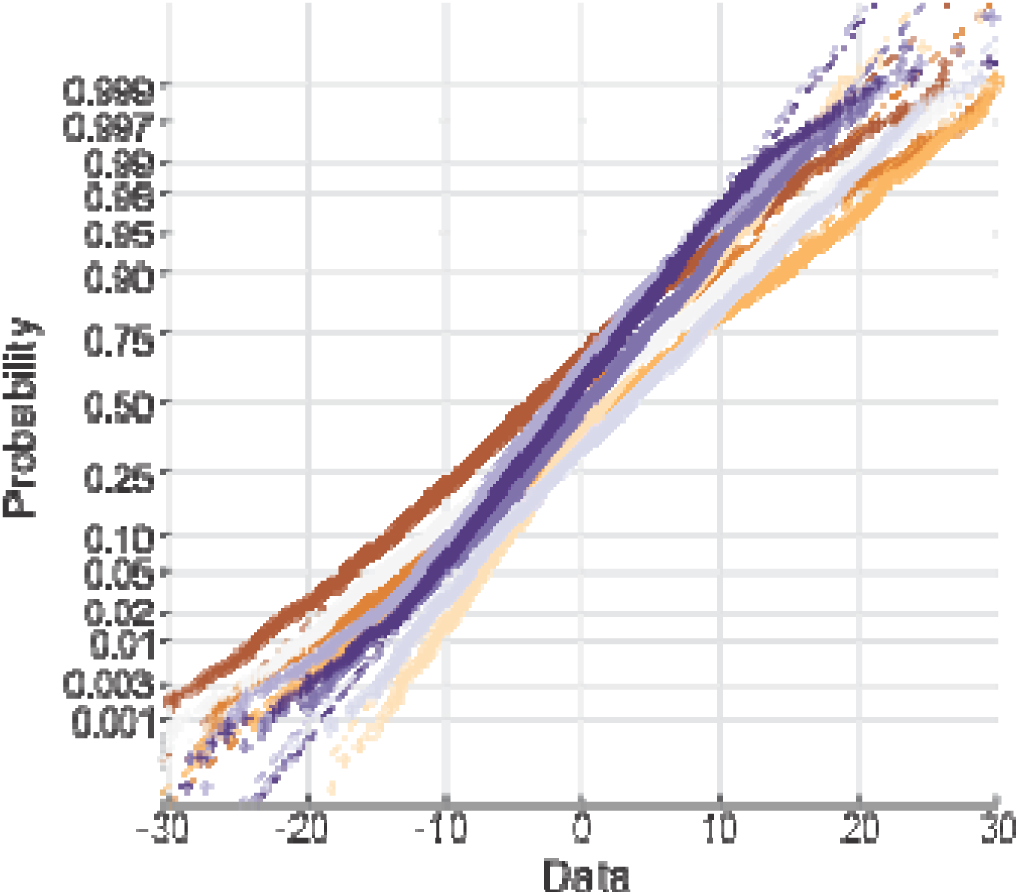
Residual analysis: normal probability plot. The probability plot is depicted for each participant, different colors. As it can be seen the residuals of our model fit compared to the participants’ data has almost a normal distribution for all the participants.

Finally, we examined if our residual has a random pattern by examining the normality of our model residual using normal probability plot, plotted using MATLAB 2016 ‘normplot.m’. Figure 11 provides the normal probability plot of our fitting for all nine participants. As it is depicted, residual values for all the participants approximately have a normal distribution which implicates that our model captures all the features in the data. More details of how our model explains the data can be found in supplementary materials.

## Discussion

We assessed the effect of neck muscle spindle noise on multi-sensory integration during a reaching task and found that applying neck load biased head angle estimation across all head roll angles resulting in systematic shifts in reach errors. We also examined the effect of head roll on reach errors and observed both an increase in movement variability and biases in reaching errors; similar to Burns & Blohm (2010). To quantitatively examine the effect of noise on reaching movements, we developed a novel 3D stochastic model of multisensory integration across reference frames. The effect of neck muscle spindle noise and head roll could be explained by a misestimation of head orientation and signal-dependent noise in the RFTs between visual and proprioceptive coordinates. The model was able to successfully reproduce the reaching patterns observed in the data providing evidence that the brain has online knowledge of full body geometry as well as the reliability associated with each signal and uses this information to plan the reach movement in a statistically optimal manner.

### Model discussion

In our model, the multisensory integration process occurs in specific reference frames; i.e. in visual and proprioceptive coordinates. Therefore, signals should be transformed into the appropriate coordinate frame before integration which is done by a series of coordinate rotations and translations. However, we do not claim that the brain performs these computations in the same explicit serial way. Alternatively, neurons could directly combine different signals across reference frames (Abedi Khoozani et al., 2016; Blohm et al., 2009; Ma et al. 2006; Beck et al., 2011), e.g. by gain modulation mechanisms. Regardless of the mechanism used, we expect very similar behavioral outcomes.

In addition, we assumed that all the distributions remain Gaussian after performing RFT processes to simplify the required Bayesian computations.However, in general, this is not necessarily correct. For example, it has been shown that noisy transformations can dramatically change the distribution of transformed signals (Alikhanian et al., 2015). Since the noise in our RFTs was small enough, the deviations from a Gaussian distribution are negligible and this approximation did not affect our model behavior dramatically. It would be interesting, though, to examine how considering the actual distribution and performing the basic Bayesian statistics (Press, 1989) will change the model behavior.

### Interpretation of observations

We suggest that neck load biased head angle estimation across all head roll angles, which resulted in systematic biases in reach error curves. Our model accounted for these shifts by assuming that neck load biases the head angle estimation toward the direction of the load. How our brain estimates the head orientation has previously been investigated. The vestibular system and especially otolith system is very important for estimating the static head orientation relative to gravitational axes (Fernandez et al., 1972; Sadeghi et al., 2007). Vingerhoets et al. (2009) demonstrated that tilted visual and vestibular cues bias the perception of visual verticality. The author showed that a Bayesian model that integrates visual and vestibular cues can capture the observed biases in verticality perception. Furthermore, muscle spindles play an important role in determining joint position sense (Goodwin et al., 1972; Scott & Loeb, 1994) compared to the other sources; e.g. tendons or cutaneous receptors (Gandevia et al., 1992 and Jones, 1994). Armstrong et al. (2008) showed that the muscles in the cervical section of the spine have a high density of muscle spindles providing an accurate representation of head position relative to the body. Therefore, head angle can be estimated from a combination of visual, vestibular and neck muscle spindle information.

We included multisensory integration of visual/vestibular, and neck muscle spindle signals in our model. Since we only modulated neck muscle information, we assumed that a combination of visual/vestibular signals is integrated with neck muscle spindle information. We were able to retrieve the relative contribution of visual/vestibular information vs. neck muscle spindle information by fitting our model to the data. We found that the contribution of neck muscle spindle information was very low (in the order of 5%) compared to visual/vestibular information.

There could be several possible explanations for observing a relatively low contribution of the neck muscle information. First, we selected the amount of the neck load in a way to apply force comparable to 30deg head tilt. However, due to the complex organization of neck muscles (Armstrong et al., 2008) we couldn’t directly measure the changes in muscles’ activity. Therefore, to accurately measure the effect of applying load on neck muscle spindle information, a detailed model of neck muscle organization would be required. Moreover, usually neck muscle information agrees with the skin receptor (i.e. Cutaneous receptor) information. In our task, however, the neck muscle information and Cutaneous receptor information are in conflict, which might be a potential reason for down-weighting neck proprioceptive information (Körding et al. 2007).

Unexpectedly, we observed that applying head roll creates larger reaching movement biases for visual targets away from the body compared to visual targets toward the body. This pattern can be captured by including the full body geometry in the RFT processes in our model. Previously, Blohm and Crawford (2007) showed that in order to accurately plan a reaching movement to visual targets, the full body geometry (both rotations and translations) has to be taken into account by the brain. Based on our model, the displacement of centers of rotation between head- and eye-centered coordinate spaces caused this asymmetry in the reaching movements.

In addition to biases, we observed that reaching movements were more variable in the straight head with neck load conditions compared to the straight head and no load condition. We considered this as evidence for neck load affecting RFTs; we assumed that the neck muscle spindles have signal-dependent noise (Scott & Loeb, 1994). Therefore, applying the neck load increases the noise in the neck muscle spindle information and consequently the sensed head orientation. This noisier sensed head angle resulted in noisier RFTs and accordingly more variable reach movements.

Surprisingly, we observed an asymmetry in the amount of variability increase by applying neck load on the right vs. left side when the head was upright. One explanation could be that since all participants were right-handed, they were more sensitive to the changes on the left side. Several imaging studies demonstrated that right-handed people have bigger left hemispheres with more neural resources associated to the right-side of the body (Bauermeister, 1987; Linkenauger et al., 2009; Linkenauger et al., 2012). Bauermeister (1987) tested the effect of handedness on perceiving verticality and showed that right-handed participants are more responsive to the right sided stimulus than to the left sided stimulus.

Since head roll with no neck load caused higher increase in movement variability compared to applying neck load while the head was upright, we expected to see a systematic modulation of movement variability by applying neck load while the head was tilted. Specifically, we expected to observe the highest amount of movement variability when the neck load and head roll were applied on the same side; e.g. 30deg CW head roll and right side neck load. The logic is that when both head roll and neck load are applied in the same direction, the neck muscle signal indicates the highest angle and due to signal dependent noise the associated variability of head angle estimations has the highest value. However, applying the load on the same side as the tilted head did not increase the movement variability significantly compared to only tilting the head.

A possible explanation for the lower effect of applying load on the same side of titled head can be relatively low contribution of neck muscle spindle information vs. visual/vestibular information during head angle estimation, provided by our model. The remarkable observation is that even though the contribution of neck muscle information is low, applying neck load still has a tangible effect on reaching movements for all head roll conditions, observed both in our data and model predictions. Another explanation is that the brain might not integrate the visual/vestibular information in this condition due to the big discrepancy between neck muscle information and visual/vestibular information (due to lack of causality; Kording et al. 2007). In our experimental design we selected the neck load value to simulate the same force as when the head is titled 30deg (head weight*sin(60deg) = 0.5*head weight). Consequently, when the head is tilted 30deg CW and the load is applied on the right side the total force on the neck muscle can be calculated as: 0.5*head weight (head tilt)+0.5*head weight (head load) = full head weight; this force is stimulating the neck muscle as if the head was tilted 90deg, which is very unlikely. Therefore, the brain might ignore the neck muscle spindle information fully.

We interpreted the observed increase in movement variability as an indication of signal dependent noise in the RFT process. However, an alternative hypothesis to the signal dependent noise is uncommon posture (Körding & Sabes 2011). According to the uncommon posture hypothesis, we might have more neuronal resources allocated to the head straight posture since we perform most of our movements with the head straight. As a result, rolling the head creates higher uncertainty due to uncommon posture independent of signal dependent noise. Even though this argument might be valid for head roll, it cannot explain the increased movement variability due to applying neck load. In other words, applying neck load while the body posture was kept unchanged still increased the movement variability which is in contrary to the uncommon posture hypothesis.

We observed that changing head roll and/or applying neck load modulated multisensory weights in both visual and proprioceptual coordinates. We validated this finding by both fitting Sober and Sabes’ (2003) model and our new full Bayesian RFTs model to the data. We found that increasing the noise in the sensed head angle estimation decreased the reliability of transformed signals, which we hypothesized is the result of the stochastic RFTs, and consequently lowered weights for transformed signals in the multisensory integration process. Therefore, we conclude that both body geometry and signal-dependent noise influence multi-sensory integration weights through stochastic RFTs.

We demonstrated that head position estimation plays a vital role in RFT processes required for reach movements. Previous studies showed that non-visual information of head position in space, i.e. from the neck muscle spindles (Proske & Gandevia, 2012) and from the vestibular system (Angelaki & Cullen 2008; Cullen 2012), decline over time providing better information about the relative changes in head position than about the absolute position. This behavior is explained based on proprioceptive drift; afferent discharges decline over time (Tsay et al., 2014) resulting in imprecise absolute estimation of the head position. We evaluated the temporal evolution of head angle estimation and its possible effect on reaching movements by dividing each block of our experiment into 4 bins (data not shown); however we found no changes in movement biases or variability. Therefore, we believe that our experiment design and timing was not appropriate to investigate changes of head angle estimation over time on reaching behaviour. This question, however, is an intriguing question and should be investigated in future experiments.

### Implications

Our findings have implications for behavioral, perceptual, electrophysiology, and modeling studies. First, we have demonstrated that both body geometry and stochasticity in RFTs modulate multisensory integration weights. It is possible that other contextual variables such as attention or prior knowledge also modulate multisensory weights and will subsequently affect both perception and action. In addition, we have shown that such modulations in multisensory weights can create asymmetrical biases in reach movements. Such unexpected biases may be prevalent in behavioral data obtained during visuomotor experiments in which participants perform the task in a robotic setup while their body is in various geometries, e.g. tilted head forward or elevated elbow. Therefore, it is important to consider that forcing specific body configurations can create unpredicted effects that are important for interpreting the behavioral data.

Our findings also suggest that the brain must have online knowledge of the statistical properties of the signals involved in multisensory integration. This could be achieved by population codes in the brain (Ma et al., 2009), which agrees with the current dominant view that the brain performs the required computations through probabilistic inferences (Pitkow & Angelaki, 2017). Alternatively, multisensory weights and the change of weights with contextual parameters could be learned (Mikula et al., 2018). Learned weights could be specially advantageous when it is difficult to estimate sensory reliability. Computational models that include required latent variables are crucial to understand the required computations. An important benefit of such models is that they can be used to generate training sets for neural networks in order to investigate potential neural mechanisms underlying probabilistic inference. Such studies will motivate appropriate electrophysiology experiments to validate/refute predications of related models.

## Supplementary materials

In this section we provide more details of how our model performs RFTs on different sensory signals (modeled as Gaussian distributions). In the manuscript, we demonstrated that the proposed model was able to replicate our behavioral data pattern (Figure 9). In this section, we use the model to provide a mechanistic explanation of the observed reach movement patterns.

Sober and Sabes (2003) demonstrated that reaching errors caused by dissociating visual and proprioceptive information can be explained by two components: MV error that is the error at the vector planning stage and INV error which is the error at the motor command generation stage. They showed that adding these two reaching errors leads to the error pattern observed in human participants. Furthermore, Burns and Blohm (2010) demonstrated that the observed up- and downward shifts in reaching error curves can be explained by RFTs; any misestimation in the sensed head angle results in an erroneous rotation of movement vector which results in up- and downward shifts in reach error curves. The logic is the same in our model for explaining the observed biases in reach error curves for the head roll condition. Similarly, the up/down - ward shifts in reach error curves for the neck load condition can be explained by erroneous RFTs; applying a neck load biases the head angle, which leads to an erroneous rotation of the movement vector, resulting in shifts of error curves.

Applying a neck load enabled us to evaluate the contribution of neck muscle spindle information to head angle estimation. To achieve this we included a Bayesian head angle estimator in our model in which the visual/vestibular and neck muscle spindle information are integrated to estimate the head angle. Applying a neck load biases the neck muscle spindle information toward the direction of the load and consequently biases the head angle estimation (equations 7 and 8). This bias in estimated head angle depends on two parameters: 1) relative neck muscle reliability compared to visual/vestibular reliability and 2) overall head angle estimation variability (similar to the variable RFTs variance in Burns and Blohm’s (2010) model)**Error! Reference source not found.**.

As explained before, in our model we estimate the head angle by integrating visual/vestibular information with neck muscle information. As a result, the overall sensed head angle variability depends on the variability of each of aforementioned information. Consider the situation in which overall head angle estimation variability is low (**Error! Reference source not found.** Figure S1 A-B). Low variability for head angle estimation resulted from high reliability for both visual/vestibular and neck muscle spindle information. Similarly, high variability of head angle estimation resulted from low reliability of both visual/vestibular and neck muscle spindle information and consequently applying neck load creates smaller biases (**Error! Reference source not found.** Figure S1 C-D). We expect that applying a neck load will create higher shifts in reach error curves for when the reliability of sensed head angle is high compared to when the reliability of sensed head angle is low, regardless of their relative contribution (compare Figure S1A vs. Figure S1D and Figure S1B vs. Figure S1C).

In addition, the amount of shifts in reach error curves depends on the relative reliability of neck muscle spindle information vs. visual/vestibular information. When the relative reliability of neck muscle information is high, the bias in reach error curves is higher compared to when its reliability is low (Figure S1B**Error! Reference source not found.** vs. Figure S1C). In our data, we observed high variability for head angle estimation as well as relatively higher contribution of visual/vestibular information compared to neck muscle spindle information (); Figure S1C**Error! Reference source not found.**.

**Figure S1.**
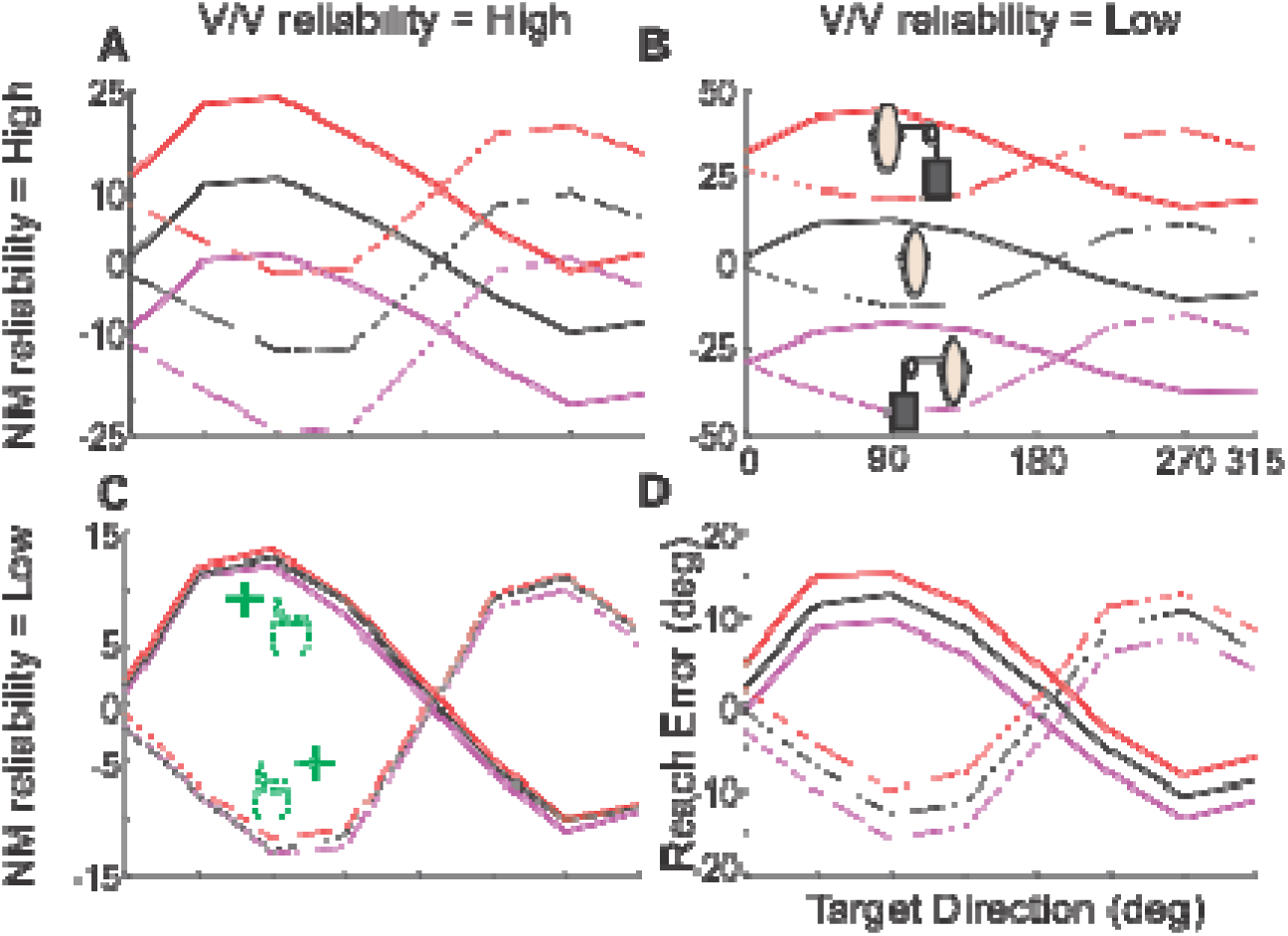
Effect of varying the reliability of neck muscle spindle signals vs. visual/vestibular signals. Head angle is estimated by combining the neck muscle spindle information with combined visual and vestibular information using the Bayesian method, therefore, the effect of applying neck load depends on two factors: 1) absolute variability of head angle estimation and 2) relative reliability of neck muscle spindle information compared to visual/vestibular information. A-B) lower absolute value for head angle estimation variability: this lower variability results from the high reliability of both visual/vestibular and neck muscle information. Therefore, the up/down-ward shifts induced due to applying neck load is higher compared with when the head angle estimation variability is high (panel C and D). In addition to the absolute head angle estimation variability, the relative reliability of neck muscle spindle vs. visual/vestibular information impacts how much applying neck load biases the reaching movement: A, C) the lower the reliability of neck muscle spindle information vs. visual/vestibular information, the lower the up/down ward shifts in reaching error curves, B, D) increasing the relative reliability of neck muscle information increases the up/down ward shifts in reaching errors by applying neck load‥

As mentioned before, at the heart of our RFT process there is a head angle estimator which enabled us to retrieve the sensed head angle based on the reach error patterns. Figure S2 demonstrates the biases in head angle estimation for all the experimental conditions. As can be seen, applying neck load biased the head angle estimation toward the applied neck load for all head angles. We performed t-test analysis and observed that all the changes in head angle estimation due to applying neck load are significant −11 < t(8) < 12, p < 0.001.

**Figure S2.**
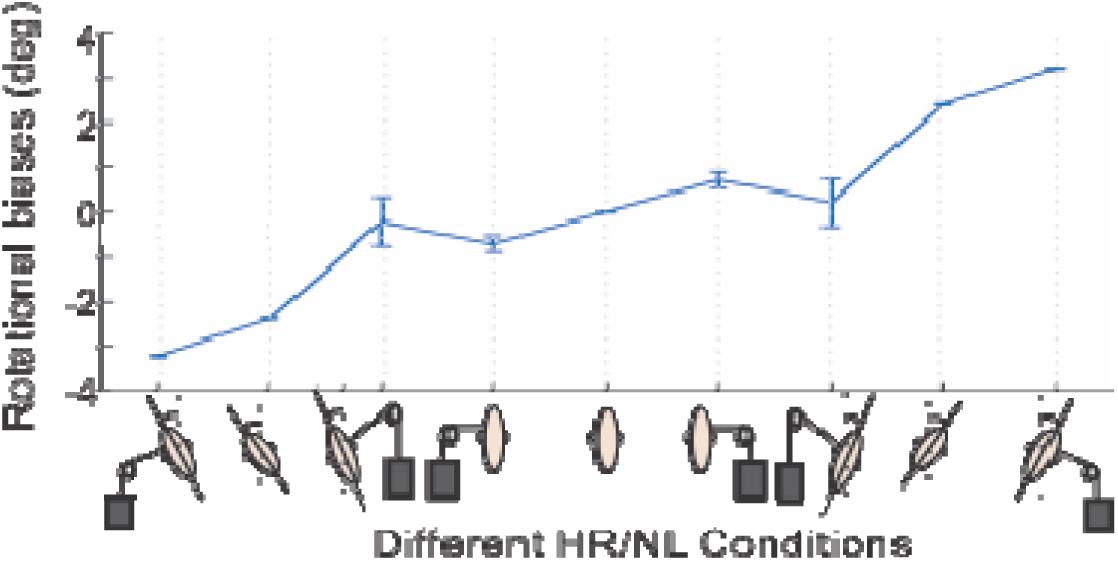
Biases in head angle estimation due to different head roll and neck load conditions. Applying neck load biased the head angle estimation toward the applied load for all head angles. Error bars are standard deviations.

In addition to up/down –ward shifts in reach error curves by applying neck load and head roll, we observed a very surprising pattern in our data: both head roll and neck load created greater biases in reaching movements when reaching to targets away from the body (45-135 deg) compared to reaching to targets toward the body (215-315 deg). This observation was surprising and to our knowledge none of the previous models (Sober & Sabes 2003 and Burns & Blohm 2010) could predict/explain this pattern.

At this point it should come as no surprise that our model explains the difference in head roll/neck load effect for different targets by stochastic RFTs processes. Blohm and Crawford (2007) demonstrated that the brain considers the full 3D body geometry to accurately plan reach movements. As mentioned in the model description, we included the 3D body geometry in our RFTs procedure: RFT processes are carried out by sequential rotations/translations between different coordinates centered on different body sections. Figure S3A demonstrates different coordinates that have been considered in our model in relation to each other. Including the 3D body geometry resulted in a displacement in the center of rotation between different coordinates and specifically in our experiment between gaze-centered and head-centered coordinates. This displacement of the center of rotation caused greater biases in reaching movements for visual targets further away from vs. closer to the body (Figure S3A). Figure S3B provides a detailed example of how the difference in the center of rotation results in an asymmetry in the movement biases induced by head roll/neck load. The first block in Figure S3B shows the actual scene in front of the participants with two targets at 90deg and 270deg. In our experiment the participants fixated their eyes on the cross and this cross was indicated as their visual information of the initial hand position as well. In this example, the hand was shifted 25cm horizontally to the right. The dotted arrows show the visual movement vector toward the targets. Box #1 demonstrates the retinal representation of targets for head roll 30deg CCW. We assumed that the torsion effect on retinal information was small and therefore ignored it. Since the head is rotated 30deg CCW, the retinal image on the back of the head is rotated 30deg CW (actual head angle) and the center of this rotation is the cross (gaze-position). In order to estimate the hand position, proprioceptive information must be transformed to the retinal coordinates and at the heart of this transformation is the head rotation based on the estimated head angle (Blue box in figure 3). In this specific example, we assumed that the head angle is overestimated by 5deg and is estimated as 35deg. In addition, since the centers of rotation for head-centered and gaze-centered coordinates are different, the transformed hand position is no longer in symmetry with the rotation in gaze-centered coordinates and displaced and biased toward the body. The next two steps in our model are multisensory integration to estimate the hand position and movement vector calculations (Box #2). As it has been shown by Sober and Sabes (2003, 2005) any transformation adds noise and therefore, visual information is more reliable in the retinal coordinates and the estimated initial hand position is biased toward the visual initial hand position and the movement vector is calculated by subtracting target position from this estimated initial hand position. This movement vector, then, is transformed into shoulder-center coordinates to be executed, employing RFTs (Box #3). We compared the transformed movement vector with the visual movement vector in Box #4 and as it can be seen the misestimation in head angle created greater biases for target away from the body (90deg) compared to the target toward the body (270deg).

**Figure S3.**
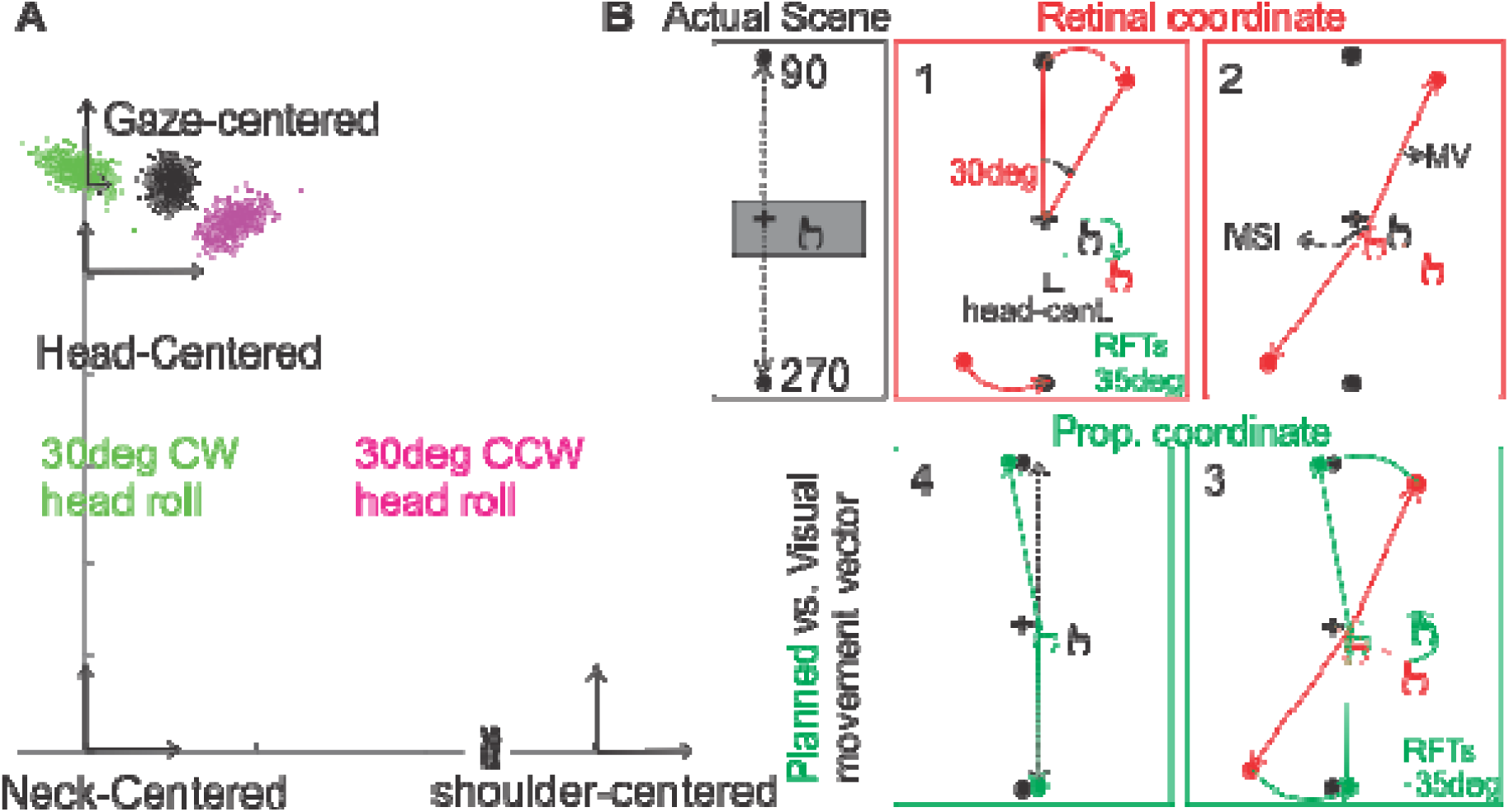
RFT processes mechanism. A) Different coordinates in our RFT module. The difference in the center of rotation between gaze-centered coordinate and head-centered coordinate resulted in an asymmetry of transformed hand position for 30deg CW vs CCW head rolls. B) A detailed example of the higher effect of stochastic RFTs on movement away from the body compared to movements toward the body for head roll 30deg CCW: Actual scene: in our experiment, participants had fixated their eyes on the center cross and the visual feedback of the hand indicated their hand on the center as well. The actual hand position is shifted to the right in this example and it is occluded, Box#1: the retinal image of the target is rotated 30 CW, we ignored the torsion effects on retinal projection. Proprioceptive hand position is transformed using our RFT module (we assumed that head roll estimation is erroneous; 35deg), Box#2: Initial hand position is estimated by combining visual information and transformed proprioceptive information of the hand. Then, the movement vector is calculated by subtracting target position from the initial hand position, Box#3: The calculated movement vector is transformed to the proprioceptive coordinate using the RFTs module, Box#4: comparing the planned movement with the movement only considering visual information. As it can be seen, the misestimation in head angle, created larger error for movement away from body vs. movement toward the body. This happened due to the offset in the center of rotations between different coordinates.

Determining how stochastic noise in RFTs modulates multi-sensory weights was one of the goals of this experiment. In figures 6 and 7, we fitted Sober and Sabes’ (2003) model to the data and demonstrated that both head roll and neck load modulates multi-sensory integration weights. Similar to Burns and Blohm (2010), we were able to retrieve multisensory integration weights from the covariance matrices. As it has been demonstrated in figure S4, RFTs dramatically change the distribution of the transformed signal and consequently the covariance matrix (Alikhanian et al., 2015). In order to account for such variations, we calculated the determinant of the covariance matrix for calculating the multi-sensory weights. Figure S4 shows visual weights in both visual (A) and proprioceptive (B) coordinates.

**Figure S4.**
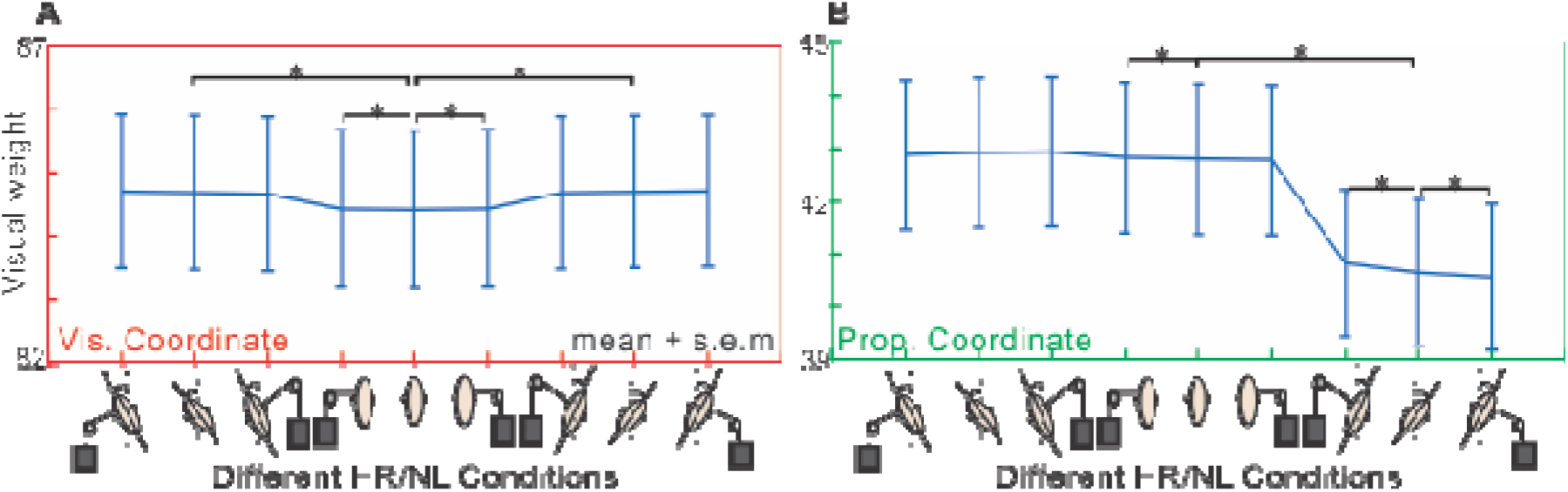
visual weights for multi-sensory integration. A) Visual weights in visual coordinate: Visual weights increase in visual coordinate due to decreased reliability of proprioceptive information caused by stochastic RFTs, B) Visual weights in proprioceptive coordinate: rolling the head 30deg CCW didn’t affect the visual weights while rolling the head 30deg CW decreased visual weights. The reason for this asymmetry is the nonlinearity in the inverse kinematic process. Error bars are standard error of the mean. The significance was tested using paired t-test (P < 0.05 is considered as a significant difference).

Visual weights were lowest for head straight and no load condition in visual coordinates and increased by rolling the head and/or applying neck load. Our paired t-test showed that this increase was significant for all head roll and neck load conditions (t(8) < −3, p < 0.05). More specifically, applying the neck load increased the visual weights in visual coordinate while the head was upright (t(8) < −3, p < 0.05) while it didn’t significantly change when the head wasn’t upright and neck load was applied (t(8) < −1, p ≈ 0.2). Applying neck load or rolling the head didn’t significantly changed visual weights in visual coordinates except for when the head rolled 30deg CW (t(8) < −18, p < 0.001) or the neck load applied to the left side (t(8) < 3, p < 0.05). Combination of head roll and neck load only modulated the visual weights when the head was rolled 30deg CW and neck load applied to the either sides (|t(8)| < 4, p < 0.05). Therefore, our data and model show that both noise in RFTs and the geometry of the body can influence multi-sensory integration in a way that is explained through changes in reliability of transformed signal by stochastic and geometrically accurate RFT processes.

